# Neural Representations of Ensemble Mean and Variance Across Visual Features

**DOI:** 10.64898/2026.07.14.738506

**Authors:** Patxi Elosegi, Seyma Takir, Ning Mei, Yaoda Xu, David Soto

**Author notes:** <u>Corresponding Author:</u> Patxi Elosegi, Department of Psychology, Yale University, New Haven, CT 06520.

## Abstract

Humans can rapidly extract summary statistics, such as the mean and variance of visual features, to efficiently represent complex visual environments despite limits in attention and working memory. For example, when viewing a field of flowers, we can perceive the average colour and overall variability of the display without individuating each flower. Although ensemble perception is central to visual cognition, two fundamental questions remain unresolved. First, it is unclear whether ensemble statistics for features represented at different levels of the visual hierarchy rely on a common neural system or on separate feature-specific systems. Second, it remains unknown whether different summary statistics, such as mean and variance, rely on shared or dissociable neural mechanisms. Here, we used fMRI and multivariate pattern analysis to examine the neural representation of ensemble mean and variance across three visual features spanning the processing hierarchy: orientation (low-level), shape (mid-level), and animacy (high-level) (N = 24; two fMRI sessions). By combining whole-brain searchlight and ROI-based approaches, we found a graded division of labour between ventral and dorsal visual pathways. Although mean and variance ensemble statistics were distributed across the visual cortex, mean decoding was stronger in ventral regions, whereas variance decoding was stronger in dorsal regions. Ensemble mean representations followed a posterior–anterior gradient within the ventral visual pathway, consistent with increasing abstraction from orientation to shape and animacy, and showed little anatomical overlap suggesting largely feature-specific. By contrast, ensemble variance was weighted toward dorsal parietal and frontoparietal regions, especially superior parietal cortex and intraparietal sulcus, decoding clusters largely overlap across features and generalized robustly across orientation, shape, and animacy. Together, these findings provide a more nuanced account of ensemble perception, showing that feature-specific and feature-independent neural codes can coexist across visual cortex and help reconcile previously conflicting evidence.

**Significance Statement:** This study shows that information about ensemble mean and variance is distributed throughout the visual cortex but weighted differently across visual pathways. Mean-related information was stronger in the ventral visual pathway and showed a largely feature-independent anatomical organization, whereas variance-related information was stronger in dorsal parietal regions and generalized across orientation, shape and animacy. Together, these findings provide clarifying neural evidence for longstanding debates about whether ensemble statistics rely on shared or dissociable mechanisms and whether ensemble representations are feature-specific or generalize across visual features. More broadly, the study offers a nuanced account of ensemble perception in which feature-specific and feature-general neural codes coexist across visual cortex, potentially reconciling previously conflicting evidence.

## 1. Introduction

Neuroscience studies of visual cognition have traditionally focused on how the brain represents isolated objects; however, objects in the real world rarely appear alone. In fact, natural scenes contain far more items than the visual system can individuate at once (i.e. navigating through a crowd or assessing the flow of traffic) but, fortunately, this clutter is not completely random. Instead, natural scenes are often filled with similar or redundant groups of objects, features, and textures. When the number of items exceeds our attentional and working memory limits, the visual system capitalizes on this redundancy by extracting a summary statistic (e.g., mean or variance) of the set of objects—known as *ensemble perception* (Alvarez, 2011). This ability allows us, for example, to effortlessly estimate the average color and diversity of flowers in a meadow without inspecting each one individually.

Ensemble representations are versatile, extending across various modalities and feature dimensions such as orientation, size, motion, animacy, and even crowd emotion (see Whitney & Yamanashi Leib, 2018; Corbett, Utochkin & Hochstein, 2023). Notably, ensemble perception appears to operate automatically, persisting under crowding (Parkes et al., 2001) minimal attention (Alvarez and Oliva, 2008; Chong and Treisman, 2005), and arguably without perceptual awareness (Elosegi et al., 2024).

However, despite the importance of ensemble perception, its neural underpinnings remain poorly understood. A central unresolved question is whether ensemble coding relies on domain-general mechanisms that operate across visual features or on feature-specific processes tailored to particular stimulus dimensions (e.g., orientation, shape, or animacy; Whitney & Yamanashi Leib, 2018; Corbett et al., 2023). Much of the evidence addressing this question comes from individual-difference approaches, which assess whether performance covaries across ensemble tasks involving different visual features (Haberman et al., 2015). Supporting a feature-independent account, several studies report robust correlations in performance across diverse visual ensemble tasks (Kacin et al., 2021; Chang & Gauthier, 2021). However, other studies found no significant correlations across different visual features, potentially suggesting independent mechanisms (Haberman et al., 2015; Yörük & Boduroglu, 2020).

Similarly unresolved is whether distinct ensemble descriptors—such as mean and variance—rely on shared or separate neural mechanisms. Behavioral evidence here is also inconsistent. For example, significant correlations between judgments of mean size and size variability (Cha et al., 2021) or mean and variance in color (Hansmann-Roth et al., 2021) suggest shared processing mechanisms. Conversely, independent processing channels have been proposed, based on the absence of correlations between judgments of mean and variance for orientation and size ensembles (Utochkin & Vostrikov, 2017; Yang et al., 2018; Khvostov & Utochkin, 2019).

Given the inherent limitations of behavioral correlations in disentangling neural mechanisms (Corbett et al., 2023), direct neuroimaging approaches are necessary. Early fMRI studies identified the parahippocampal place area (PPA)—linked to scene and texture perception (Epstein & Kanwisher, 1998; Cant & Goodale, 2007)—as a possible locus for encoding ensemble statistics (Cant & Xu, 2012, 2015, 2017). While subsequent research has further supported the role of the PPA extracting the “gist” of a scene (Park & Park, 2017), accumulating evidence suggests ensemble representations are distributed across broader cortical networks. For instance, Im et al. (2017) reported a double dissociation, demonstrating dorsal parietal activation during the perception of mean ensemble emotion, in contrast to ventral stream activation during individual emotion recognition. Although concerns have been raised regarding methodological confounds in this study (Cant & Xu, 2020), converging evidence consistently suggests that ensemble codes extend beyond PPA. Tark et al. (2021) showed that average orientation information can be reconstructed in early visual areas, with additional frontoparietal engagement when observers focus attention on ensemble information. These findings suggest a complex, possibly hierarchical, and distributed neural basis for ensemble perception (Corbett et al., 2023).

The methodological heterogeneity of previous neuroimaging studies—including adaptation paradigms, univariate contrasts and encoding approaches, alongside substantial variation in stimulus complexity and task demands—has hindered a coherent account of the neural basis of ensemble perception. Critically, most neuroimaging work has focused exclusively on ensemble mean, leaving other fundamental statistical descriptors, such as variance, unexplored. As a result, it remains unclear whether ensemble representations generalize across visual features, and whether different summary statistics rely on shared or dissociable neural mechanisms.

Here, we address these gaps using a comprehensive functional MRI paradigm in which ensemble mean and variance were independently manipulated across different visual features of varying complexity. The neural representations underlying the different summary statistics were characterized using whole-brain multivariate searchlight analyses, coupled with region-of-interest decoding to determine whether ensemble mean and variance rely on overlapping or distinct cortical systems across statistical descriptors (i.e., mean and variance) and levels of visual complexity (i.e., orientation, shape, and animacy).

## 2. Methods

### 2.1. Participants

Following informed consent, twenty-five participants (7 males, 18 females; age range, 19–36 years; right-handed) were recruited for the study in exchange for monetary compensation at a rate of €10 per hour of experiment. One participant withdrew half way through the experiment, resulting in a final sample of twenty-four participants who completed the two fMRI sessions of the study. All participants had normal or corrected-to-normal visual acuity and reported no history of psychiatric or neurological conditions. The study adhered to the Declaration of Helsinki (2008) and was approved by the Basque Center on Cognition, Brain and Language (BCBL) Research Ethics Board. Our sample size is larger than most previous fMRI studies on ensemble perception which were based on standard mass-univariate analyses (Cant & Xu, 2012, 2015, 2017; Tark et al., 2021). Furthermore, each participant completed two fMRI sessions across two different days in order to boost the sensitivity of our decoding analyses.

### 2.2. Stimulus Generation

To examine the feature specificity of ensemble representations across visual domains, participants viewed three types of ensembles constructed from visual features spanning the visual processing hierarchy, from low-level oriented gratings, mid-level shapes (i.e., spiky–stubby), and high level animacy (i.e., animate–inanimate objects). Ensemble displays consisted of twelve objects arranged in a 4 columns x 3 rows grid (see Figure 1-A), centered around fixation. Low-level image properties such as contrast, luminance, and object size were controlled across stimulus conditions to minimize confounds using SHINEToolbox (Willenbockel et al., 2010) and in-house python scripts.

**Figure 1.**
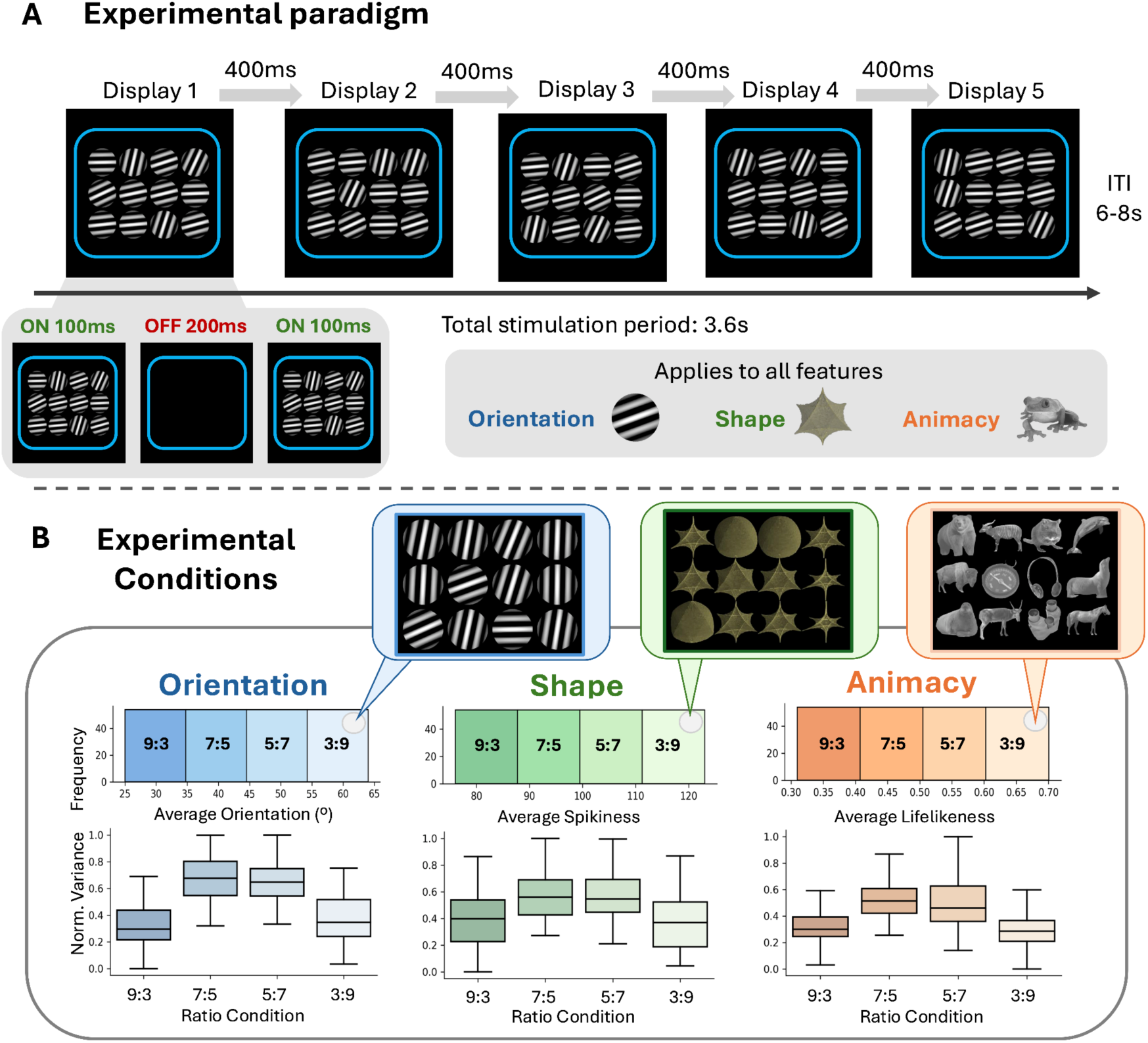
Overview of Experimental Procedure and Conditions. **A)** *Experimental paradigm:* Ensembles were presented using a miniblock design. Within each trial, the same ensemble was presented across five successive displays, with object locations randomized on each presentation. Each display was shown twice (100 ms on, 200 ms off), and successive displays were separated by a 400 ms blank interval. Following the fifth display, a jittered inter-trial interval (6–8 s) preceded the next trial. A blue frame and fixation cross remained visible throughout the trial, and each trial began with a brief fixation cue signaling stimulus onset. **B)** *Experimental conditions:* Histograms provide a visual summary of the continuous uniform distributions binarized across different ratio conditions for every visual feature. The boxplots below show the ensemble variance separately for low-variance ratio conditions (9:3 and 3:9) and high-variance ratio conditions (7:5 and 5:7). To facilitate comparison across visual features, variance values were normalized within each feature dimension and plotted on a common scale. For orientation, the x-axis of the histogram represents average grating orientation, with ratios indicating the proportion of steep to shallow gratings in each ensemble (e.g., 3 steep : 9 shallow). For shape, the x-axis of the histogram represents the spikiness continuum, with ratios indicating the number of stubby versus spiky objects (e.g., 3 stubby : 9 spiky). For animacy, the x-axis of the histogram depicts the lifelikeness continuum, with ratios reflecting the proportion of animate to inanimate objects (e.g., 3 inanimate : 9 animate). For each visual feature, an example ensemble from the 3:9 condition is depicted on top of the histograms.

To examine whether different summary statistical descriptors of ensembles are represented within shared neural substrates, we focused on two core summary statistical descriptors: the ensemble mean and the ensemble variance. The ensemble mean—defined as the average feature value across items in a set (e.g., mean orientation, spikiness, or animacy)—has been the most extensively studied metric in the ensemble perception literature (e.g., Ariely, 2001, Whitney & Yamanashi Leib, 2018). In contrast, ensemble variance—capturing the degree of variability among features within a set—represents a distinct type of summary statistic that has received comparatively little empirical attention (Morgan et al., 2008; Lau & Brady, 2018), despite its potential relevance for perceptual stability and scene understanding (Corbett et al., 2023).

During the generation of ensemble stimuli, single objects were sampled to ensure that mean ensemble features varied continuously along a uniform distribution (see upper histograms in Fig. 1-B). This distribution was partitioned into four equally sized, non-overlapping but immediately adjacent bins, each corresponding to specific ratio conditions: 9:3, 7:5, 5:7, and 3:9. These ratios defined the proportion of items from the two distinct classes used in each ensemble type. For example, in the animacy condition, a 9:3 ratio represented nine animate and three inanimate objects, while a 3:9 ratio represented the inverse (see below for condition-specific details). Accordingly, the first half of the distribution (9:3 and 7:5) reflected a predominance of one category (i.e., animate), whereas the second half (5:7 and 3:9) reflected a predominance of the other (i.e., inanimate).

Importantly, the symmetrical structure of the mirror ratios (i.e., 9:3 and 3:9, 7:5 and 5:7) enabled to orthogonally manipulate the ensemble mean and variance information. The 9:3 and 3:9 conditions had relatively low variance with most of the ensemble elements from one category (see bottom boxplots in Fig. 1-B). In contrast, the 7:5 and 5:7 conditions had relatively high variance with more equal contributions from both categories. This design ensured that variance and category dominance could be independently studied.

For each stimulus condition (i.e., orientation, shape, animacy), 216 unique ensembles were generated (54 ensembles per ratio condition) and during the experiment, each unique ensemble was presented only once.

#### 2.2.1. Orientation Ensembles

Each orientation ensemble contained Gabor patterns in different orientations (spatial frequency = 0.62 cycles per degree). An initial pool of 50 Gabor patches was generated by dichotomizing the orientation continuum into two categories: 25 *steep* and 25 *shallow* Gabors. Steep Gabors were sampled from orientations ranging from 65° to 89°, whereas shallow Gabors were sampled from orientations ranging from 1° to 25°. This stimulus pool was subsequently used to construct the ensemble ratio conditions.

Specifically, for each ratio, 10,000 ensemble combinations were randomly sampled to generate a distribution of mean ensemble orientations. Based on these distributions, non-overlapping ranges of average orientations were defined for each ratio. Specifically, for the 9:3 ratio (9 shallow vs. 3 steep), the average orientation ranged from 25° to 34°; for 7:5, it ranged from 35° to 44°; for 5:7, from 45° to 54°; and for 3:9, from 55° to 65°. Thus, mean ensemble orientations across ratios spanned from 25° to 65°. Fifty-four ensembles were then generated for a given ratio condition in the defined mean ensemble orientation range, resulting in a total of 216 orientation ensembles across the four ratio conditions.

#### 2.2.2. Shape Ensembles

Shapes varied along a spikiness dimension ranging from stubby (smooth, rounded contours) to spiky (contours with pronounced protrusions and sharp vertices). This dimension has been identified as a principal axis of object representation in both human and non-human primate visual cortex (Bao et al., 2020, Xu & Chun, 2025).

In our study, spiky and stubby objects were parameterized by rendering geometrical patterns in three dimensions using the p-norm function, which includes a parameter *p* that controls the level of regularization. For instance, L1 and L2 regularizations, commonly used in machine learning, correspond to *p* = 1 and *p* = 2 respectively (see <u>Equation 1</u>.).

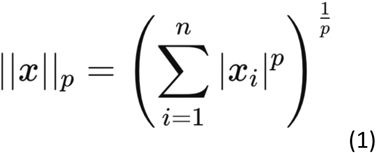

Apart from regularization, the *p* parameter also defines the geometrical characteristics of the p-norm function. Values between 0 and 1 produce star-shaped or concave forms, while values between 1 and 2 transition from diamond to near-spherical shapes. By systematically varying *p* from 0.4 to 1.6, we generated a continuous spectrum of shapes parametrized along a "spikiness" dimension.

The continuum of spikiness was categorized into two classes: *spiky* shapes (ranging from *p* = 0.4 to *p* = 0.89) and *stubby* shapes (ranging from *p* = 1.1 to *p* = 1.6). Following the same procedure used to generate the orientation ensemble, here, for each ratio, 10,000 ensemble combinations were randomly sampled and the resulting spikiness distributions were used to define for each ratio non-overlapping, equal-size ranges of spikiness. Specifically, the spikiness ranges were 0.76–0.88 for the 9:3 ratio (9 stubby vs. 3 spiky), 0.89–1.0 for the 7:5 ratio (7 stubby vs. 5 spiky), 1.1–1.12 for the 5:7 ratio (5 stubby vs. 5 spiky), and 1.13–1.24 for the 3:9 ratio (3 stubby vs. 9 spiky). For each ratio, 54 unique ensembles were generated, totaling 216 unique shape ensembles).

#### 2.2.3. Animacy Ensembles

Each animacy ensemble contained a mixture of living and non-living objects. Unlike orientation and shape which have a continuous distribution of feature values, animacy is inherently a categorical feature (i.e., living versus non-living). Unlike orientation or shape, which occupy well-defined continuous feature spaces, animacy is conventionally treated as a categorical distinction. However, recent evidence suggests that ‘lifelikeness’ varies along a continuous neural dimension within both living and non-living categories (Sha et al., 2015; Leib et al., 2016; Thorat et al., 2019). This inherent variability allowed us to manipulate animacy magnitude, analogous to parametric variations in orientation or spikiness.

Accordingly, we sought to estimate a continuous measure of *lifelikeness* for each item, leveraging a computational approach. Specifically, 120 grayscale images of living and non-living objects were drawn from the Konkle Lab stimulus set (https://konklab.fas.harvard.edu/#, Long et al., 2018). Each image was processed through an ImageNet pre-trained DenseNet-169 convolutional neural network (CNN). DenseNet-169 has been shown to produce representations similar to those found in the human inferotemporal (IT) cortex (Schrimpf et al., 2018), thus providing a good approximation to object representation in the ventral visual pathway.

To compute lifelikeness scores, we first generated a *prototypical living* representation by averaging the activation patterns of the penultimate layer activations of the CNN across all living stimuli. The lifelikeness of each object was then quantified by calculating the cosine similarity between the layer activations of each object and the prototypical living vector. Consequently, objects closer to the prototypical living representation were assigned higher lifelikeness scores.

It is important to acknowledge that, even if CNNs are the current best model of IT cortex, they can only explain about 60% of its variance (Yamins et al., 2014; Schrimpf et al., 2020). We therefore validated our CNN-derived lifelikeness estimates against independent human behavioral data obtained from two independent ensemble-perception experiments (N=60). Human-derived and CNN-derived lifelikeness estimates were highly correlated at both the individual-object and ensemble levels (*r*_s_ = 0.88–0.95), indicating substantial correspondence between the two measures. Details of this validation analysis are reported in the Supplementary Materials 1.

Following the procedure used for orientation and shape ensembles, we generated 10,000 random combinations of objects for each ratio condition. The resulting distributions were used to define non-overlapping lifelikeness ranges for each ratio. Specifically, the ranges of lifelikeness were: 0.3–0.4 (where 0 means non-living and 1 mean living) for the 9:3 ratio (9 non-living vs. 3 living objects), 0.4–0.5 for the 7:5 ratio (7 non-living vs. 5 living objects), 0.5–0.6 for the 5:7 ratio (5 non-living vs. 7 living objects), and 0.6–0.7 for the 3:9 ratio (3 non-living vs. 9 living objects). A total of 216 unique animacy ensembles were generated.

### 2.3. Study Overview

MRI data were collected across two experimental sessions separated by no more than two weeks. All stimulus presentation scripts were generated in Python 3.9 using the OpenSesame library (Mathôt et al., 2012). Each session lasted approximately 1.5 hours, including participant preparation and breaks between runs.

### 2.4. Procedure

Ensembles were presented using a miniblock design. Within each trial, the same ensemble was shown across five successive displays, with object locations randomized on each presentation (Fig. 1A). This manipulation preserved ensemble composition while reducing correlations in low-level visual input across displays. To facilitate the formation of a stable ensemble representation, each display was presented twice in an ON–OFF–ON sequence (100 ms ON, 200 ms OFF, 100 ms ON). Although cumulative stimulus exposure was limited to 200 ms, the intervening blank interval extended processing time to 400 ms while remaining below the latency typically required for voluntary saccadic eye movements (Carpenter, 1988). Consecutive displays were separated by a 400 ms blank interval, except following the fifth display, yielding a total stimulation period of 3.6 s. Each trial was followed by a jittered inter-trial interval (ITI) of 6–8 s.

Stimuli subtending 7.96° × 7.04° of visual angle were presented on a dark-gray background within a blue frame (Figure 1-A). The frame delineated the ensemble region and signaled trial onset by briefly flickering 500 ms before stimulus presentation. Participants were instructed to maintain central fixation throughout the experiment and to attend to the ensemble as a whole rather than to individual items, thereby encouraging holistic ensemble processing.

Across the two scanning sessions, participants completed 720 trials. Within each session, they performed three consecutive runs of each stimulus condition (animacy, shape, and orientation). The order of condition triplets was counterbalanced across participants such that each condition appeared equally often at the beginning, middle, and end of a session. Furthermore, no participant experienced the same triplet order across the two sessions.

Each run comprised 40 trials, including 36 valid trials and 4 catch trials. Valid trials were evenly distributed across the four ratio conditions (9 trials per condition). Participants were encouraged to take short breaks between runs and were informed of the upcoming stimulus condition during these intervals.

Catch trials were included to ensure sustained attention to the ensemble displays. Participants performed an oddball detection task, responding with their right index finger only when they detected a catch trial within a 1.5-s response window following stimulus offset. Catch trials initially contained a 10:2 ratio of items and, during the third or fourth display, reversed to a 2:10 ratio (or vice versa). Twenty-four unique catch stimuli were generated for each stimulus condition, with each catch trial presented only once during the experiment. To reduce predictability, a minimum of four valid trials separated any two catch trials.

At the beginning of each session, participants completed extensive training both inside and outside the scanner with performance feedback. They were instructed to adopt a conservative response criterion and to respond only when confident that a catch trial had occurred. This instruction was intended to minimize false alarms, as both catch trials and false-alarm trials were excluded from the primary analyses. To ensure adequate task engagement, participants were required to achieve at least 70% catch-trial detection accuracy across stimulus conditions. All participants met this criterion, and none were excluded.

### 2.5. MRI Data Acquisition and Preprocessing

fMRI data were collected with a 3T 3-T Siemens Magnetom Prisma-fit scanner and a 64-channel head coil at the Basque Center on Brain, Cognition and Language (BCBL). In each fMRI session, a multi-band gradient-echo echo-planar imaging sequence with an acceleration factor of 6, resolution of 2.4×2.4×2.4 mm^3^, repetition time of 850 ms, echo time of 35 ms and bandwidth of 2,582 Hz/Px was used to obtain 550 three-dimensional (3D) volumes of the whole brain (66 slices; field of view 210 mm). For each observer, one high-resolution T1-weighted structural image was also collected. Visual stimuli were projected on an MRI-compatible out-of-bore screen using a projector placed in the room adjacent to the MRI room, and viewed through a small mirror mounted on the head coil by the participants while they lied inside the scanner. The head coil was also equipped with a microphone, allowing verbal communication between the participant and experimenters between the runs.

All MRI data were converted from DICOM to NIfTI format using MRIConvert (http://lcni.uoregon.edu/downloads/mriconvert). The converted data were subsequently processed using FEAT (fMRI Expert Analysis Tool) from the FMRIB Software Library (FSL suite; v6.0.3). To ensure steady-state magnetization, the first eight volumes of each functional run were discarded. Brain extraction was performed using FSL’s Brain Extraction Tool (BET; v2.1), effectively removing non-brain tissue (Smith, 2002). We then applied the Automatic Removal of Motion Artifacts (AROMA) method to identify and eliminate motion-induced signal fluctuations (Pruim et al., 2015). The data were spatially smoothed with a Gaussian kernel of 3 mm full width at half maximum and subjected to a high-pass filter with a cutoff frequency of 60 seconds to remove low-frequency drifts. All scans were co-registered to a reference volume derived from the first run of the first session to ensure alignment across the data sets.

Following the preprocessing of the MRI data, we utilized outputs generated by OpenSesame during the experimental task to label the relevant scans. Each preprocessed scan was annotated with attributes such as ratio condition (e.g., 9:3, 7:5, 5:7, or 3:9), predominant class (e.g., predominantly spiky vs stubby), continuous ensemble features (i.e., average orientation, spikiness, or lifelikeness) and variance condition (i.e., high vs low). Data from both sessions were combined after linear detrending of each voxel’s time series. To account for hemodynamic lag, we generated each trial example for decoding analysis by averaging the pre-processed scans four to seven seconds following the onset of each of the five ensemble variants, which resulted in the averaging of 7–8 volumes per trial (Pereira et al. 2009). To prevent differences in overall response magnitude from driving classification, the multivoxel activity pattern at each time point was z-scored within each run and condition (i.e., 9:3, 7:5, 5:7, 3:9) across all voxels within the pattern (e.g., Vaziri-Pashkam & Xu, 2017).

### 2.6. Multivariate Pattern Analysis

Multivariate pattern analysis (MVPA) provides a means of characterizing the information encoded in distributed patterns of neural activity and, in the present context, of determining how ensemble mean and variance are represented across visual features. By comparing the cortical distribution of decoding effects across orientation, shape and animacy, we could assess whether ensemble representations are supported by common, anatomically centralized regions or distributed cortical systems, and whether this organization differs between statistical descriptors. For ensemble variance, we additionally tested whether representations generalized across features using cross-decoding. Successful cross-decoding would indicate that variance information is encoded through a shared representational mechanism across visual features. An analogous analysis was not feasible for ensemble mean because the classification labels used for orientation, shape and animacy lack a principled correspondence; any mapping between them would therefore be arbitrary.

#### 2.6.1 Whole-Brain Searchlight

MVPA embedded in a whole-brain searchlight algorithm (Kriegeskorte et al., 2006) was employed to identify cortical regions encoding ensemble-level information. A spherical searchlight with a 6 mm radius was used. All analyses were implemented in Python with the Scikit-learn (Pedregosa et al., 2011) and Nilearn (Abraham et al., 2014) libraries, and were performed on a within-subject basis in each participant’s native space.

To assess statistical significance, the results from each participant were averaged across cross-validation folds, and the resulting 3D brain maps were normalized to standard space. The 3D maps of all 24 participants were then concatenated into a single 4D file, which was subsequently used as input for the *randomise* tool in FSL (Smith et al., 2004) for non-parametric permutation statistical testing using the Threshold-Free Cluster Enhancement (TFCE) method to identify clusters of significance above chance classification/regression without the need to set an arbitrary threshold for voxel-level statistics (Nichols and Holmes et al., 2002). In the following, we detailed the specific procedure followed for each MVPA analysis.

##### 2.6.1.1. Ensemble Mean Decoding

To identify the mean ensemble feature independently for orientation, shape, and animacy ensembles, we employed a binary classification analysis to decode the predominant class (e.g., [9:3, 7:5] vs. [5:7, 3:9]) independently for each stimulus condition. The rationale is that if the multivoxel pattern within a searchlight window captures the mean ensemble feature, a classifier distinguishing the predominant class could effectively position a decision boundary at the midpoint of the average feature distribution.

We employed Linear Support Vector Classifiers (SVC), trained and tested using leave-one-run-out (LORO) cross-validation with six folds per stimulus condition. This cross-validation scheme minimizes the influence of within-run correlations between training and test sets, providing a more conservative estimate of decoding performance.

For every classification analysis, default parameters for SCV provided by Sklearn (C=1) were applied without additional hyperparameter tuning, ensuring a direct comparison across all classification analyses. Receiver operating characteristic area under the curve (ROC–AUC) was used to quantify classification performance, as it can provide a more sensitive measure than raw accuracy in machine-learning classification tasks (Bradley, 1997; Fahrenfort et al., 2018).

In parallel, we conducted a regularized ridge regression to localize brain regions that predicted the mean ensemble value along the continuous feature dimension. This regression analysis served as a sanity check, ensuring that the predominant class decoding approach captured representations of the underlying average feature, not simply categorical distinctions. Within each searchlight and for every stimulus type, we fit an L2-regularized multivariate linear model using LORO cross-validation. For each training split, a nested 5-fold grid search was conducted to optimize the alpha parameter of ridge regression, which controls regularization strength and model complexity. Following Conwell et al. (2024), the grid search evaluated alpha values at 10, 10^2^, 10^4^, 10^5^, and 10^7^. Cross-validated explained variance (cvR^2^) was used as a metric of models’ prediction performance.

##### 2.6.1.2. Ensemble Variance Decoding

To identify feature-specific representations of ensemble variance, we performed a separate binary classification to discriminate between low-variance (i.e., more homogeneous) from high-variance (i.e., more heterogeneous) ensembles (e.g., [3:9, 9:3] vs. [5:7, 7:5]). This analysis was also run independently for each stimulus condition. Following the same approach as for the predominant class decoding, linear SVCs were used for this analysis, embedded within a LORO cross-validation protocol.

##### 2.6.1.3 Ensemble Variance Cross-Decoding Analysis

To test to what extent ensemble representations are feature-independent, SVCs were trained to decode a particular ensemble statistic (i.e., variance) using data from two stimulus conditions (e.g., orientation and shape) and then tested on a third condition (e.g., animacy). This was done iteratively, ensuring that all combinations of stimulus conditions were part of the training and test sets. Training across two conditions increased the amount and diversity of the training data, encouraging the classifier to capture feature-general variance representations while reducing overfitting to any single stimulus dimension.

For robust generalization, within each combination of training and testing partitions (i.e., train on orientation+shape and test on animacy; train on orientation+animacy test on shape; train on shape+animacy and test on orientation), we performed 20 iterations of a Stratified Shuffle Split (SSS) cross-validation protocol. In each iteration, we randomly sampled 80% of the training data (e.g, orientation+shape), ensuring an equal number of items from each stimulus condition and the classifier was subsequently tested on the held-out stimulus condition (e.g., animacy). This repeated subsampling procedure assessed the robustness of cross-decoding to variation in the training sample.

Although training and testing were conducted across stimulus conditions, results are reported separately for each test condition. For example, results for shape reflect classifier performance when trained on orientation and animacy and tested on shape.

#### 2.6.2. Region of Interest Multivariate Patterns Analysis

Although searchlight analyses provide a comprehensive, whole-brain assessment of ensemble representations, group-level inference can be limited by inter-individual anatomical variability. In addition, the restricted spatial extent of the searchlight spheres may reduce sensitivity to distributed representations spanning larger cortical territories. These limitations are particularly relevant for our goal of comparing dorsal and ventral involvement in ensemble perception. Accordingly, to directly assess differences between dorsal and ventral visual pathways in ensemble coding, we complemented the searchlight approach with a region-of-interest (ROI)–based multivariate pattern analysis.

Our ROIs encompassed key visual and parietal regions, including pericalcarine cortex, lingual gyrus, lateral occipital cortex, fusiform gyrus, inferior temporal gyrus, inferior parietal gyrus, and superior parietal gyrus (see Fig. 6-A). Cortical parcellation was performed on high-resolution structural images using FreeSurfer (recon-all, version 6.0.0). The resulting anatomical ROI masks were registered to native functional space using seven-degree-of-freedom linear registration in FSL FLIRT, binarized, and used to extract multivoxel response patterns for subsequent decoding analyses.

To assess pathway-level differences, we additionally constructed composite ROIs by combining anatomically defined regions. The ventral composite mask included pericalcarine, lingual, lateral occipital, fusiform, and inferior temporal regions (including the parahippocampal gyrus and adjacent temporal areas), whereas the dorsal composite mask comprised inferior and superior parietal regions (see Fig. 6-A).

All decoding and cross-decoding analyses were conducted in native BOLD space for each participant within each ROI and composite ROI, using the same classifier, cross-validation scheme, and parameter settings as in the searchlight analyses.

Statistical significance of decoding performance was evaluated at both the within-participant and group levels. Within participants, permutation testing was used: label vectors were randomly shuffled and classification was repeated 10,000 times to generate an empirical null distribution. P-values were computed as the probability of observing the true decoding performance under this distribution. At the group level, mean decoding performance was compared against the mean of the empirical chance distribution using paired-samples t-tests across participants. All p-values were corrected for multiple comparisons across ROIs, stimulus conditions (orientation, shape, animacy), and analysis types (within- and cross-decoding), separately for variance and predominant-class analyses, using Bonferroni correction.

## 3. Results

### 3.1. Behavioral Results

Average oddball detection accuracies were above 90% across conditions. As shown in Figure 2-C, no participants were excluded for low performance on the oddball task, as all participants exceeded the 70% accuracy threshold across conditions (e.g., even participant 11 who showed the lowest performance scored 71% in the animacy condition). This indicates that participants maintained attention to ensemble information throughout the experiment. Notably, most errors were misses rather than false alarms, thereby, confirming that participants effectively applied the recommended conservative response criterion (Figure 2-D).

**Figure 2.**
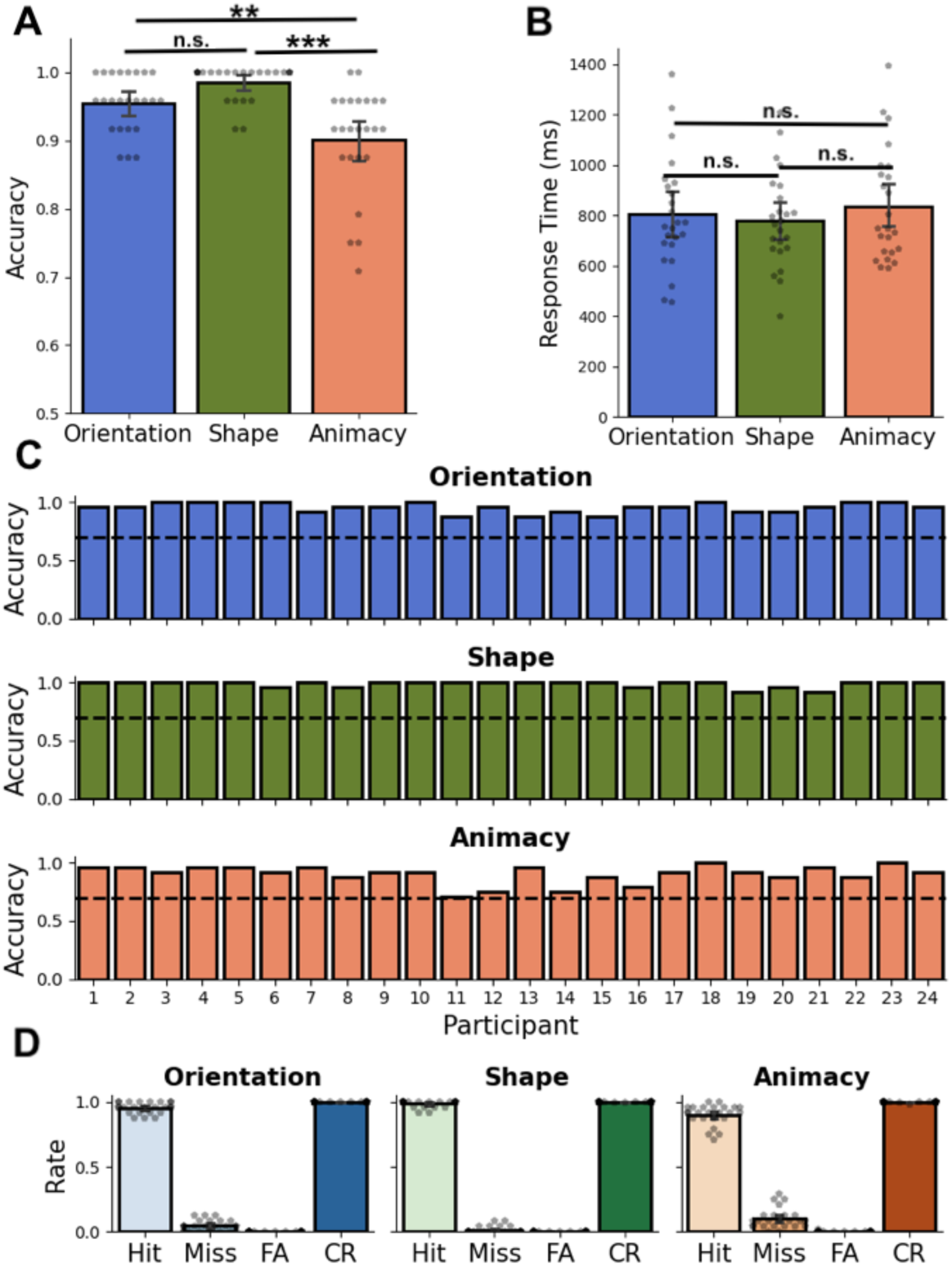
Behavioral results from the oddball detection task. Participants were required to detect four catch trials embedded within each run of the main experiment. As opposed to valid trials (e.g., 3:9, 5:7, 7:5, and 9:3), catch trials featured an extreme 10:2 ratio that reversed the predominant class to 2:10 (see Section 2.4). **A)** Displays group-level accuracy for each stimulus condition. **B)** Shows group-level response times for hit trials (i.e., successfully detected catch trials). **C)** Illustrates the accuracy of each participant across stimulus conditions, with a dashed line indicating the 70% accuracy threshold used as an exclusion criterion. **D)** Depicts the rates of trial types based on signal detection theory. The hit rate is calculated as the number of hits divided by the total number of signal-present trials (hits + misses). The miss rate is computed similarly, dividing the number of misses by signal-present trials. The false alarm (FA) rate is determined by dividing the number of false alarms by the total number of signal-absent trials (FAs + correct rejections). Lastly, the correct rejection (CR) rate is calculated by dividing the number of correct rejections by the total signal-absent trials. (n.s.: p > .5, ∗ : *p* < .05, ∗∗ : *p* < .01, ∗∗∗: *p* < .001.).

Even at the overall high-performance levels, the stimulus condition significantly influenced accuracy, as indicated by a repeated-measures ANOVA (F(1.52, 35.11) = 18.55, *p* < .001, η² = .44; Figure 2-A). Post-hoc tests revealed that all visual feature conditions differed significantly in accuracy except orientation and shape (orientation vs. shape: t(23) = −2.21, *p* = .09, d = −0.58; orientation vs. animacy: t(23) = 3.80, *p* = .002, d = 1.0; shape vs. animacy: t(23) = 6.02, *p* < .001, d = 1.58). However, differences in accuracy did not extend to reaction times since no main effect of visual feature on reaction times for correct detections (i.e., hits) was found (F(2, 46) = 1.78, *p* = .17, η² = .07; Figure 2-B).

In summary, these behavioral results confirm that participants correctly followed task instructions and remained attentive to ensemble information across all three stimulus conditions throughout the experiment.

### 3.2. Whole-brain Searchlight Multivariate Pattern Analysis Results

To determine whether neural representations of visual ensembles are feature-specific or feature-general, we conducted whole-brain multivariate searchlight analyses using a 6-mm-radius searchlight (Kriegeskorte et al., 2006). Within each visual feature, we decoded ensemble mean by discriminating the predominant class of the display (for example, [3:9, 5:7] versus [7:5, 9:3]) and ensemble variance by discriminating distributions with low versus high dispersion (for example, [3:9, 9:3] versus [5:7, 7:5]). We further tested whether variance representations generalized across features using cross-decoding, training classifiers on two feature domains and testing them on the held-out domain. An equivalent analysis was not possible for ensemble mean because predominant-class labels lacked a principled correspondence across orientation, shape and animacy (see Methods 2.6).

#### 3.2.1. Mean ensemble features are encoded along the ventral visual pathway in a feature-dependent manner

Figure 3 reveals that ensemble representations are organized along a posterior–anterior gradient within the ventral visual pathway (VVP). In early visual cortex, multivoxel activity patterns reliably discriminated whether orientation ensembles were predominantly steep (i.e., vertical, >45°) or shallow (i.e., horizontal, <45°), with decoding performance significantly above chance (Figure 3-A).

**Figure 3.**
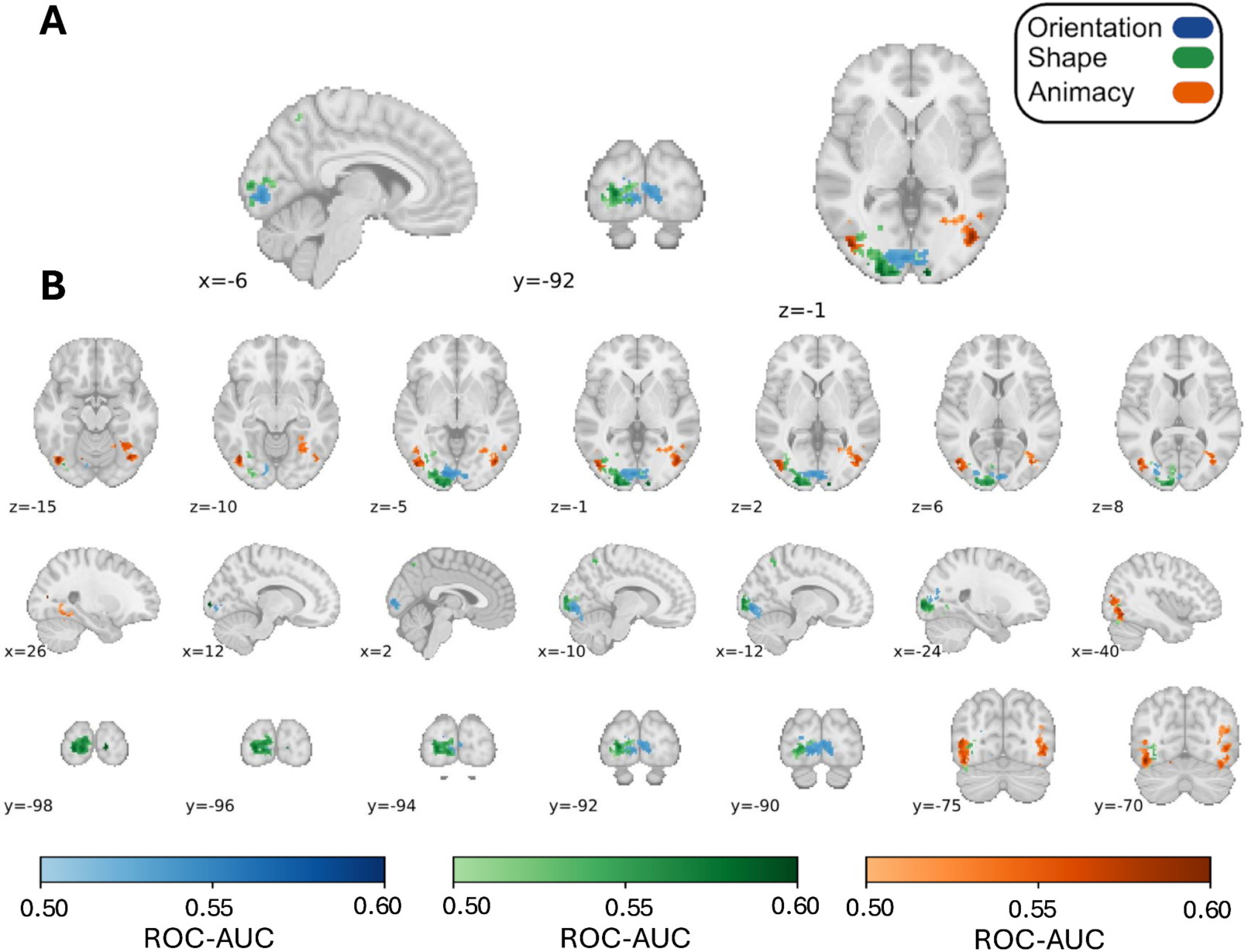
Whole-brain Searchlight Predominant Class Decoding Results Across Conditions. The heatmap indicates clusters where ROC-AUC scores were statistically significantly greater than chance (i.e., ROC-AUC = 0.5), determined using threshold-free cluster enhancement (*p* < .05) for each stimulus condition independently. **A**) highlights clusters with the highest ROC-AUC values across axial, sagittal, and coronal planes. **B**) provides detailed views of cluster locations across the brain, with rows ordered by axial, sagittal, and coronal planes, respectively. All figures are presented in the neurological view (i.e., the left side of the brain appears on the left side of the image).

Shape ensembles (predominantly spiky versus stubby) were decoded in partially overlapping but more spatially extended clusters that reached into lateral occipital cortex, consistent with evidence that this region supports shape-based object representations beyond low-level image features (e.g., Kourtzi & Kanwisher, 2001). More anteriorly, decoding of the predominant animacy category (predominantly animate vs inanimate) was observed in lateral occipitotemporal cortex and more anteriorly around the fusiform gyrus (Fig. 3B).

To verify that predominant-class decoding reflected the underlying continuous ensemble mean rather than only categorical differences, we also used ridge regression to predict the mean feature value within each searchlight. These analyses yielded convergent effects for orientation and animacy (Supplementary Fig. 3), but not for shape. The absence of significant shape regression may reflect the stricter demands of continuous linear prediction under many-to-one mapping, whereby distinct neural population states can give rise to similar ensemble estimates and therefore weaken the relationship between multivoxel activity and the mean feature value (Westlin et al., 2023).

Together, these findings suggest the mean of visual ensembles may be encoded along a hierarchical gradient within the VVP, such that progressively more anterior regions represent increasingly abstract ensemble features. This pattern is reminiscent of the canonical hierarchical organization described for single-object vision (e.g., Kravitz et al., 2013) and is broadly consistent with accounts proposing that the ventral pathway supports a graded progression from simple to more complex feature representations along an occipitotemporal axis (Posani et al., 2025; Badwal et al., 2025). Our findings extend this principle to ensemble perception, suggesting that summary statistics of visual scenes may be encoded within a similar feature-dependent representational gradient.

#### 3.2.2. Variance information is encoded in the dorsal visual pathway in a feature-independent manner

Ensemble variance decoding analysis revealed a markedly different pattern of results with significant clusters of above-chance decoding predominantly along the dorsal visual pathway (DVP) and frontal territories (see Figure 4). In particular, multivoxel patterns in bilateral superior parietal regions along the intraparietal sulcus (IPS) discriminated with significantly above-chance ROC-AUC homogenous (i.e., 9:3, 3:9) from heterogenous (i.e., 7:5, 5:7) ensembles (See Figure 5). Anterior higher-order regions, especially the right middle frontal gyrus, were also involved in encoding ensemble variance information (Figure 4-A,B, x=46, z=30). Notably, these areas showed considerable overlap across stimulus conditions, with larger parietal clusters for orientation ensembles compared to other conditions.

**Figure 4.**
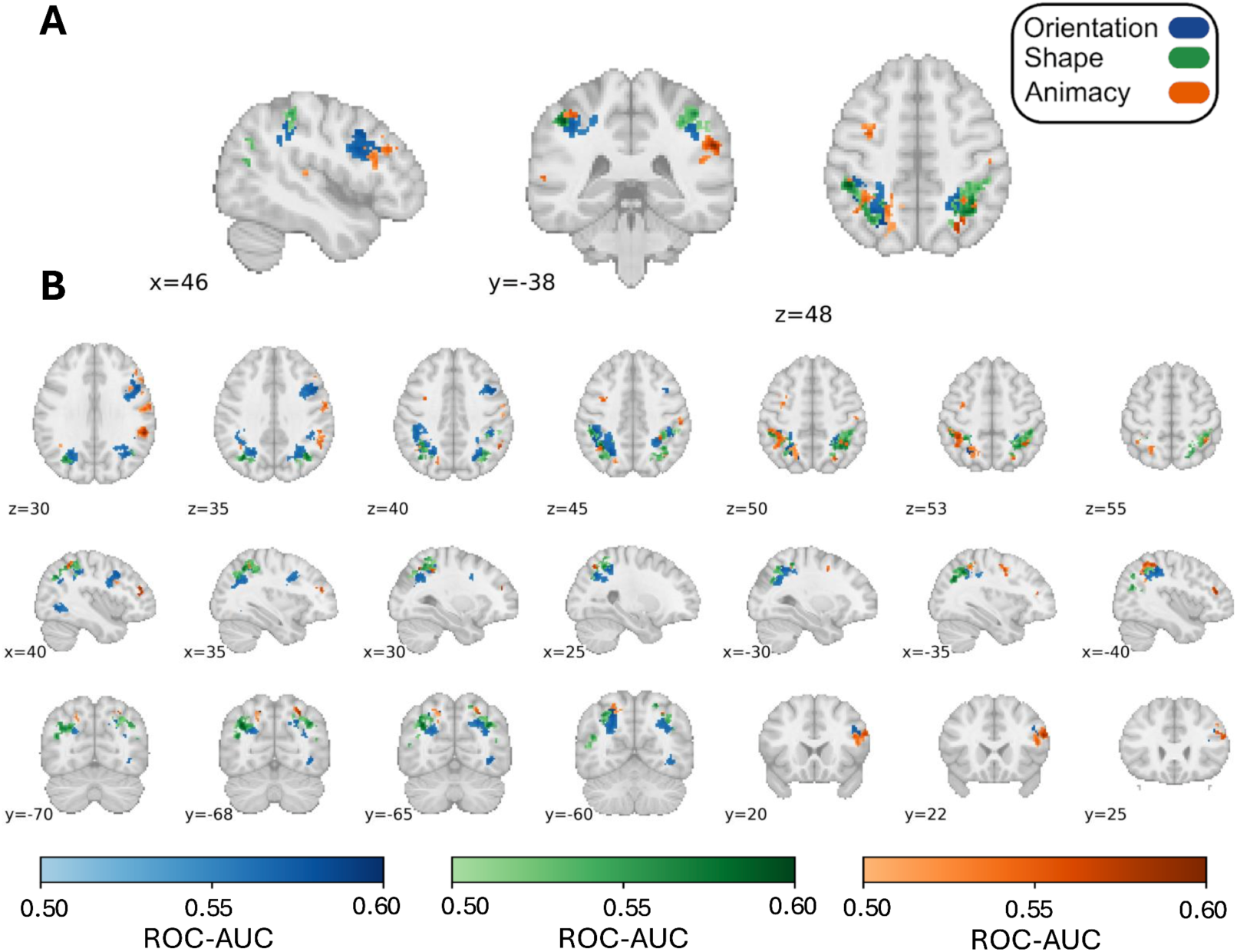
Whole-brain Searchlight Variance Decoding Results Across Conditions. The heatmap indicates clusters where ROC-AUC scores were statistically significantly greater than chance (i.e., ROC-AUC = 0.5), determined using threshold-free cluster enhancement (p < .05) for each stimulus condition independently. **A)** highlights clusters with the highest ROC-AUC values across axial, sagittal, and coronal planes. **B)** provides detailed views of cluster locations across the brain, with rows ordered by axial, sagittal, and coronal planes, respectively. All figures are presented in the neurological view (i.e., the left side of the brain appears on the left side of the image).

**Figure 5.**
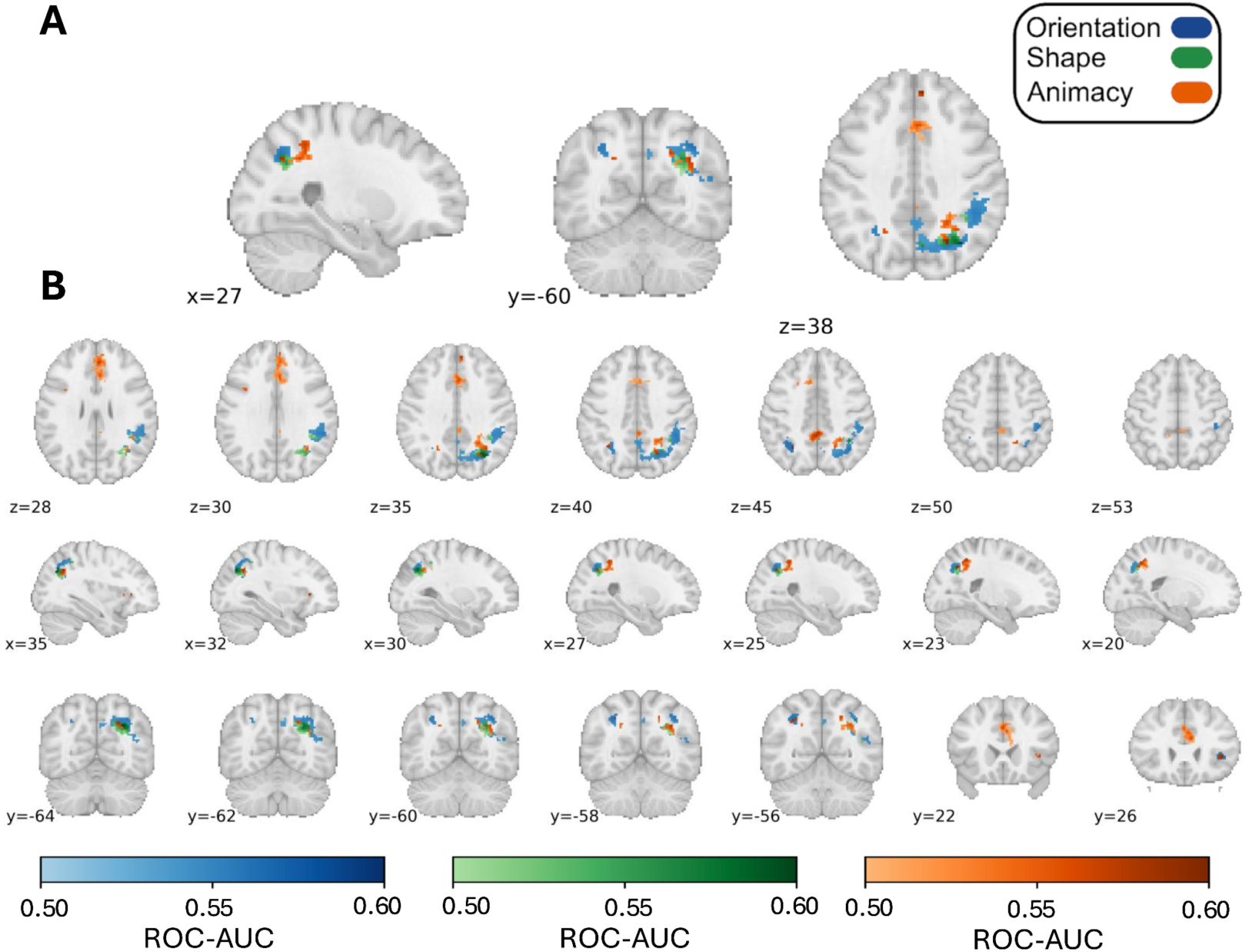
Whole-brain Searchlight Cross-Decoding of Ensemble Variance. Classifiers were trained on two conditions to predict the ensemble variance (i.e., [3:9, 9:3] vs [5:7, 7:5]) of the ensemble and tested on a held-out condition. The heatmap indicates clusters where ROC-AUC scores were statistically significantly greater than chance (i.e., ROC-AUC = 0.5), determined using threshold-free cluster enhancement (p < .05). **A**) highlights clusters with the highest ROC-AUC values across axial, sagittal, and coronal planes. **B**) provides detailed views of cluster locations across the brain, with rows ordered by axial, sagittal, and coronal planes, respectively. All figures are presented in the neurological view (i.e., the left side of the brain appears on the left side of the image).

Although ensemble variance decoding was predominantly observed along parietal and frontal regions, we also found above-chance variance decoding of orientation ensembles in the right lateral occipital cortex along the VVP (Figure 4-B, x=40).

Critically, variance cross-decoding identified a distributed network whose activity patterns generalized across stimulus features (see Figure 5). The dominant effects were centered in right superior parietal cortex, encompassing the superior parietal gyrus and posterior-to-middle intraparietal sulcus (IPS), as well as a few additional clusters in the medial frontal cortex. Together, these findings suggest that feature-invariant coding of ensemble variance relies primarily on dorsal parietal cortex, with more sparse involvement of frontal regions.

### 3.3. Region of Interest Multivariate Pattern Analysis Results

Although searchlight analyses provided a whole-brain characterization of ensemble representations of mean and variance, they are inherently most sensitive to spatially limited and anatomically consistent effects across subjects. The relatively small searchlight window (6 mm³) reduces sensitivity to representations distributed across broader cortical territories, and group-level inference further depends on sufficient spatial alignment across individuals. Consequently, searchlight analyses likely capture only the most spatially localized and topographically consistent components of ensemble coding.

We therefore complemented the searchlight MVPA with ROI-based analyses that reproduced all classification analyses described above, focusing on ventral and dorsal visual cortices to obtain a more complete characterization of the neural basis of ensemble representations. This approach also enabled a more direct test of the ventral–dorsal dissociation suggested by the searchlight findings and provided greater sensitivity to pathway differences in spatially distributed codes.

For all statistical tests of decoding performance against chance, see Supplementary Tables 1 and 2. These tables report mean group-level performance, group-level t-tests, Bayes factors quantifying evidence for above-chance performance, and the proportion of subjects showing above-chance performance according to within-subjects permutation tests. Direct comparisons between dorsal and ventral ROIs, as well as interaction with ensemble statistics (i.e., predominant-class and variance decoding), are reported in Section 3.3.2.

#### 3.3.1. Ensemble mean information is distributed across ventral and dorsal visual pathways in a largely feature-dependent manner

Predominant-class decoding at the ROI level largely mirrored the whole-brain searchlight results, while revealing a more spatially distributed pattern of ensemble representations (Fig. 6-B). To formally assess these effects, we fitted two independent linear mixed-effects models, one for ventral ROIs and one for dorsal ROIs, each including ROI and stimulus condition as fixed factors and subject as a random intercept.

**Figure 6.**
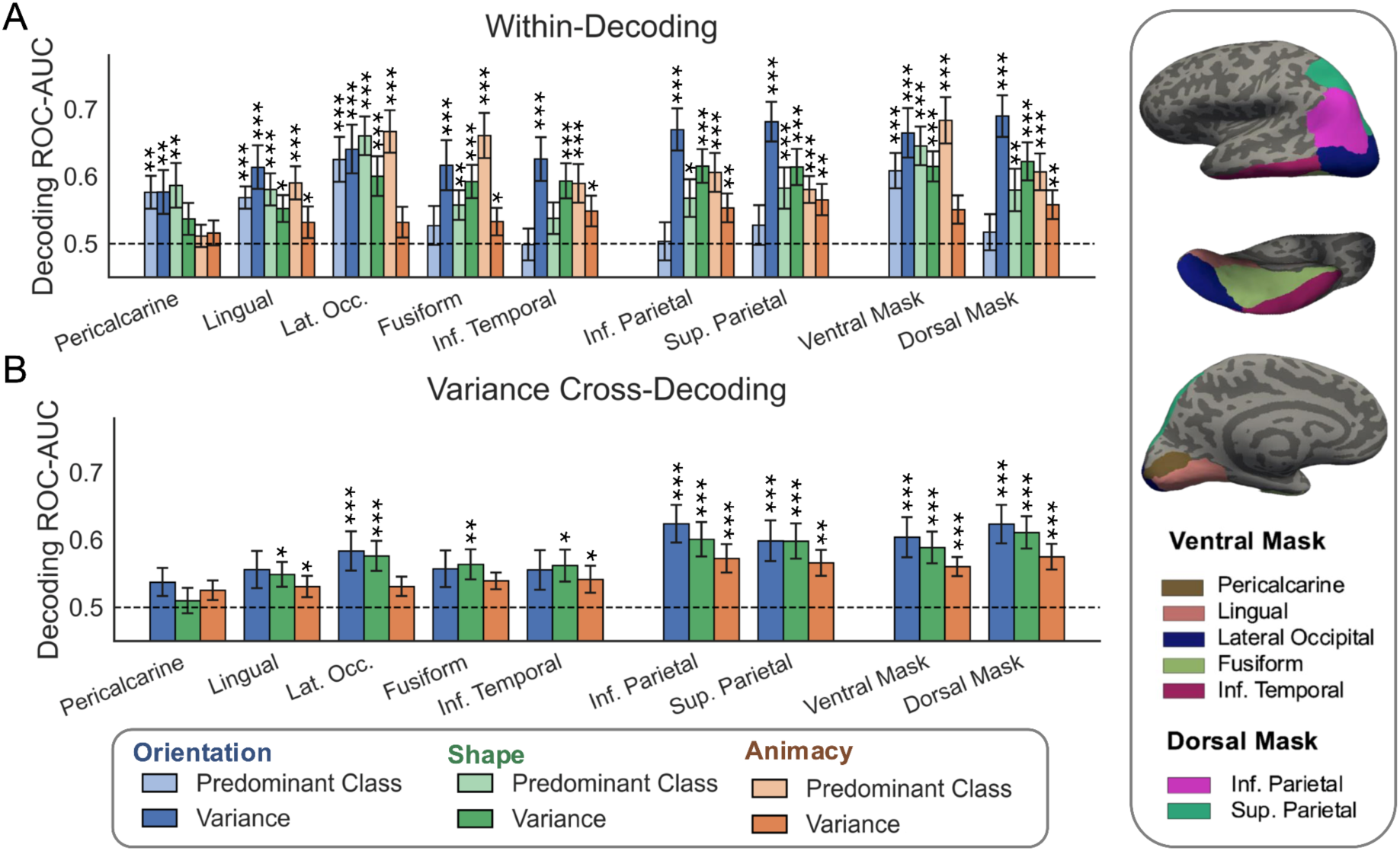
ROI-level decoding within and cross-decoding results. **A)** Within-condition decoding of predominant class (i.e,ensemble mean), and ensemble variance across ROIs, shown separately for orientation, shape, and animacy. Light bars indicate predominant-class decoding and dark bars indicate variance decoding. **B)** Cross-decoding of ensemble variance across stimulus dimensions in the same ROIs. In both panels, individual ROIs are arranged from ventral to dorsal, followed by the combined ventral and dorsal masks. The surface renderings show the ROIs on the left hemisphere. The ventral visual pathway (VVP) mask comprises pericalcarine cortex, lingual gyrus, lateral occipital cortex, fusiform gyrus, and inferior temporal gyrus; the dorsal visual pathway (DVP) mask comprises inferior and superior parietal cortex. The dashed line indicates chance-level performance (ROC–AUC = 0.50). Asterisks indicate above-chance decoding (P < 0.05, P < 0.01, P < 0.001; group-level paired-samples t-tests, Bonferroni-corrected). Error bars denote 95% confidence intervals.

In the ventral model, we observed a robust ROI × stimulus condition interaction (*F*(8,322)=11.68, *p*<.001), indicating that the spatial distribution of decoding depended on feature type. Model estimates confirmed that, in early visual cortex (pericalcarine), decoding was higher for orientation (β=0.065, SE=0.017, *t*=3.81, *p*<.001) and shape (β=0.075, SE=0.017, *t*=4.42, *p*<.001) than for animacy. This pattern progressively reversed along the ventral stream, with more anterior regions showing enhanced animacy decoding, particularly in fusiform cortex (β=0.149, SE=0.017, *t*=8.77, *p*<.001), consistent with a posterior–anterior gradient in representational complexity.

Across stimulus conditions, the strongest predominant-class decoding was consistently observed in the lateral occipital cortex. Post hoc contrasts comparing lateral occipital cortex against the mean of the remaining ventral ROIs confirmed a robust advantage across all conditions, including orientation (*t*(322)=6.14, *p*<0.001, Δ=0.083, 95% CI [0.056, 0.109]), shape (*t*(322)=7.03, *p*<0.001, Δ=0.095, 95% CI [0.068, 0.121]), and animacy (*t*(322)=5.84, *p*<.001, Δ=0.079, 95% CI [0.052, 0.105]). However, a direct paired-samples comparison between lateral occipital and fusiform cortex for animacy revealed no significant difference (*t*(23)=0.49, *p=* .63, *d*=0.09), indicating that peak animacy decoding was comparably expressed in both regions.

In contrast, the dorsal model revealed no main effect of ROI (*F*(1,115)=0.25, *p*=.62) or ROI × stimulus condition interaction (*F*(2,115)=2.63, *p*=.076). Consistent with this, inferior and superior parietal cortex did not differ overall (β=−0.025, *t*=−1.56, *p*=.123), although a modest interaction indicated that the reduction in orientation decoding relative to animacy was less pronounced in superior parietal cortex (β=0.050, *t*=2.16, *p*=.033), with a similar but non-significant trend for shape (β=0.041, *t*=1.76, *p*=.082).

#### 3.3.2. Variance ensemble information is distributed across ventral and dorsal visual pathways in a largely feature-independent manner

Variance decoding at the ROI level broadly replicated the searchlight decoding and cross-decoding results, confirming a dominant role of the dorsal visual pathway while also revealing additional, albeit weaker, involvement of ventral regions (Fig. 6-C). Variance decoding was analysed using two separate linear mixed-effects models for ventral and dorsal ROIs, each including ROI and stimulus condition as fixed factors and subject as a random intercept.

In the ventral model, we observed significant main effects of ROI (*F*(4,322)=8.07, *p*<.001) and stimulus condition (*F*(2,322)=58.11, *p*<.001), but no ROI × stimulus condition interaction (*F*(8,322)=1.10, *p*=.361). Model estimates indicated that variance decoding was higher for orientation (β=0.061, SE=0.017, *t*=3.56, *p*<.001) relative to animacy, whereas shape did not differ reliably (β=0.021, *t*=1.23, *p*=.221). Differences across ventral ROIs were modest, with no region showing a pronounced advantage. This relatively uniform pattern of above-chance variance decoding contrasts with the searchlight results, which revealed only limited ventral involvement—primarily confined to the lateral occipital cortex for orientation ensembles. This discrepancy suggests that variance-related signals in the ventral visual pathway are spatially distributed rather than locally concentrated, with searchlight analyses capturing only local peaks of decoding.

In dorsal regions, ROI results closely matched the searchlight findings. The linear mixed-effects model revealed a strong main effect of stimulus condition (*F*(2,115)=56.34, *p<.001*), but no main effect of ROI (*F*(1,115)=0.69, *p*=.409) and no ROI × stimulus condition interaction (*F*(2,115)=0.25, *p*=.779). Model estimates confirmed higher variance decoding for orientation (β=0.117, SE=0.015, *t*=7.53, *p*<.001) and shape (β=0.062, *t*=4.03, *p<.001*) relative to animacy, with no reliable differences between inferior and superior parietal cortex (β=0.012, *t*=0.79, *p*=.432).

Consistent with the searchlight analysis, ROI analyses identified the most robust feature-invariant representations of ensemble variance within the dorsal parietal cortex (Fig. 6B). Both inferior and superior parietal cortex exhibited reliable cross-decoding across all stimulus conditions. Importantly, ROI-level analyses also revealed cross-decoding effects in ventral visual regions, including the lingual gyrus for shape and animacy, the lateral occipital cortex for orientation and shape, the fusiform gyrus for shape, and the inferior temporal gyrus for shape and animacy. Together, these findings indicate that feature-invariant representations of ensemble variance are most robustly expressed in the dorsal parietal cortex, while also being detectable in a subset of ventral visual regions.

#### 3.3.3. Differential contributions of ventral and dorsal pathways to average and variance ensemble representations

Finally, we leveraged decoding within dorsal and ventral conglomerate masks to directly test whether decoding performance varied as a function of cortical pathway (dorsal vs. ventral), ensemble statistical descriptor (mean vs. variance), and their interaction. A linear mixed-effects model with random intercepts for participant and stimulus type revealed a significant interaction between pathway and statistical descriptor (*F*(1,259)=26.61, *p*<.001), indicating that the relative advantage of each statistical descriptor depended on cortical pathway. Specifically, in the dorsal pathway, variance decoding exceeded predominant-class decoding (ROC-AUC=0.623 vs. 0.568), whereas in the ventral pathway, on average predominant-class decoding outperformed variance decoding (ROC-AUC=0.646 vs. 0.610). There was also a main effect of pathway (*F*(1,259)=13.47, *p*<.001), reflecting overall higher decoding performance in ventral relative to dorsal regions, whereas the main effect of ensemble summary statistical descriptor was not significant (*F*(1,259)=1.27, *p*=0.261), consistent with the interaction.

To further examine whether these effects varied across visual features, we fitted an additional linear mixed-effects model testing the interaction between stimulus type (animacy, orientation, shape) and ensemble statistical descriptor, with random intercepts for participant and pathway. This analysis revealed a robust interaction (*F*(2,258)=59.52, *p*<.001), indicating that the relative contribution of predominant-class and variance decoding differed across stimulus types. Estimated marginal means showed that for animacy, predominant-class decoding yielded higher performance than variance decoding (ROC-AUC=0.645 vs. 0.554), whereas for orientation, variance decoding markedly outperformed predominant-class decoding (ROC-AUC=0.677 vs. 0.563). For shape, decoding performance was comparable across the two ensemble statistics (ROC-AUC=0.619 vs. 0.612). Neither the main effect of stimulus type (*F*(2,258)=2.57, *p*=.079) nor the main effect of ensemble statistics (*F*(1,258)=1.67, *p*=.197,) reached significance, consistent with the presence of the interaction.

Together, ROI analyses confirm that predominant-class (mean) ensemble information is primarily encoded along the VVP, following a posterior–anterior gradient of representational complexity, whereas variance decoding is predominantly supported by the DVP. At the same time, the ROI approach provides a more nuanced view of ensemble encoding, revealing that both ensemble mean and variance are distributed across dorsal and ventral regions rather than being strictly segregated between pathways.

### 4. Discussion

The present study sought to characterize the neural mechanisms of ensemble perception, specifically to assess whether ensemble representations diverge across visual features (orientation, shape and animacy), summary statistical descriptors (mean and variance). To address this, we combined whole-brain searchlight analyses with ROI-based decoding, allowing us to capture both spatially localized and distributed representations of ensemble statistics across different visual features.

Whole-brain searchlight analyses revealed a weighted division of labour between ventral and dorsal visual pathways. Predominant-class information was primarily encoded along the VVP, following a posterior–anterior gradient in which increasingly abstract visual features—orientation, shape, and animacy—were represented in progressively more anterior regions. Cross-decoding analyses showed that these representations were largely feature-specific. By contrast, ensemble variance was encoded primarily along the DVP, particularly in superior parietal cortex and intraparietal sulcus, with additional extension into frontal regions. In addition, variance representations generalized across visual features, indicating that the neural representation of variance is shared to some degree across different visual features.

ROI-based analyses refined this picture by showing that the ventral–dorsal division of labour was not absolute. Instead, both predominant-class and variance-related information were distributed across the visual cortex, but with different relative weights. A significant interaction between pathway and ensemble statistics showed that predominant-class decoding was stronger in the VVP, whereas variance decoding was stronger in the DVP. This suggests in line with prior behavioral evidence (Utochkin & Vostrikov, 2017; Yang et al., 2018; Khvostov & Utochkin, 2019) that ensemble mean and variance representations may rely on at least partially different neural mechanisms. Further supporting this claim, cross-decoding analysis showed that while variance representations generalized robustly across visual features, ensemble mean representations were largely feature-dependent.

The distinction between searchlight and ROI results further suggests that searchlight analyses identify clusters of ensemble coding in regions that are more anatomically consistent across participants, whereas ROI analyses capture broader distributed information that may vary in its precise topography. This interpretation is consistent with recent work demonstrating that searchlight approaches are inherently biased toward detecting localized representations and may underestimate more distributed coding schemes (Cox et al., 2021).

Together, these findings inform two central debates in the ensemble perception literature (Corbett et al., 2023): whether different ensemble statistics rely on common or distinct neural representations, and whether ensemble representations are feature-specific or generalize across visual domains. First, our results suggest that ensemble mean and variance are supported by partially distinct representational architectures. Although both statistics were represented broadly across visual cortex, ensemble mean was weighted toward the ventral visual pathway, whereas ensemble variance was weighted toward the dorsal visual pathway as indicated by the summary statistic x pathway interaction. Second, the degree of feature invariance depended on the summary statistic being represented. Whereas ensemble variance representations generalized robustly across orientation, shape, and animacy, ensemble mean representations remained largely feature-specific, supporting previous studies that found no correlation in averaging errors across different visual features (Haberman et al., 2015; Yörük & Boduroglu, 2020).

Similarly, these findings also help reconcile inconsistencies across previous neuroimaging studies of ensemble perception. Specifically, the frontoparietal encoding of abstract ratio information aligns with findings from Im et al. (2017), whereas the ventral encoding of average feature information is more consistent with Tark et al. (2021) and, to some extent, Cant and Xu (2011–2017). This suggests that apparent discrepancies across prior studies may partly reflect differences in the ensemble statistic being measured: different neural pathways may preferentially encode different forms of summary information.

With respect to ensemble mean perception, our results align closely with the findings reported by Tark et al. (2021), showing that average orientation information in visual ensembles is represented in the early visual cortex. However, our study extends these findings by demonstrating that neural ensemble mean encoding is not restricted to low-level visual features, but also encompasses more complex properties such as shape and animacy. Moreover, ensemble mean representations appear to unfold hierarchically along the ventral visual pathway, with increasingly abstract ensemble features engaging progressively more anterior temporo-occipital regions. Specifically, average animacy information was represented in more anterior ventral and inferior temporal regions than average orientation and shape information, which were represented more strongly in posterior visual areas.

In that sense, this pattern should be interpreted as a gradient in representational prominence rather than a sharp functional dissociation. Such a graded organization is consistent with influential models of the VVP, which propose that visual representations become progressively more abstract along the posterior–anterior axis while remaining distributed across overlapping cortical territories (Haxby et al., 2001; Kravitz et al., 2013; Grill-Spector & Weiner, 2014, Posani et al., 2025; Badwal et al., 2025). This parallelism suggests that ensemble mean representations may similarly share the same feature-extraction mechanisms that support single-object perception. This interpretation is consistent with recent behavioral and computational evidence showing that ensemble judgments can be predicted from the same feature-space representations used to encode individual objects (Robinson & Brady, 2023), suggesting that ensemble and object perception may rely on a common representational substrate rather than distinct perceptual mechanisms.

However, ensemble mean representations may not be limited to early visual cortex and object-selective regions. Influential fMRI adaptation studies by Cant and Xu (2012–2017) showed that more anterior ventral regions, particularly the PPA, are sensitive to summary statistical information extracted from visual ensembles. The PPA is known to respond to a variety of statistical image properties, including texture, spatial frequency, and spatial layout (Cant & Goodale, 2007; Berman, Golomb, & Walther, 2017; Park & Park, 2017), but is most strongly associated with the processing of natural scenes (Epstein & Kanwisher, 1998; Cant & Xu, 2015). Given that our stimuli consisted of isolated visual features and object ensembles rather than naturalistic scenes, it is perhaps unsurprising that we did not observe significant decoding in the PPA. One possibility is that the PPA supports the integration of summary statistics across multiple visual dimensions within scene representations, whereas the regions identified here encode the average of specific visual features or object categories. Under this account, ensemble representations may be organized hierarchically throughout the VVP, ranging from feature-specific averages in occipito-temporal cortex to increasingly abstract and integrative summary representations of naturalistic visual environments in scene-selective cortex. Future work will be needed to determine how these different forms of ensemble coding are related and clarify the role of the PPA in ensemble perception.

Beyond the VVP, representations of both ensemble mean and variance extended into the DVP. Previous studies have similarly implicated frontoparietal networks in ensemble perception. Im et al. (2017) reported increased activation in the IPS, superior parietal lobule, and frontal cortex during ensemble processing relative to single-object viewing. However, these findings have been questioned because such contrasts may reflect differences in numerosity and attentional demands rather than ensemble coding itself (Cant & Xu, 2020). Consistent with this interpretation, Tark et al. (2021) found that ensemble orientation information could only be recovered from frontoparietal regions when ensemble statistics became task-relevant.

Attentional factors may likewise have amplified ensemble variance representations in the DVP. In particular, low-variance ensembles (3:9 and 9:3) more closely resembled the catch trials (2:10 and 10:2) than did high-variance ensembles (5:7 and 7:5), potentially increasing their attentional salience and the recruitment of parietal regions that prioritize and maintain behaviourally relevant visual information (Bettencourt and Xu, 2016; Jeong and Xu, 2016; Xu, 2018). Several features of the design and analysis nevertheless argue against a purely attentional account. Variance was not cued before stimulus onset, participants were not intentionally oriented to process and select the different levels of variance – i.e. high- and low-variance trials required no differential behavioral response–, and each multivoxel pattern was normalized across voxels and within-condition, reducing sensitivity to uniform pattern-wide differences in response amplitude that would be associated with attentional modulation (Vaziri-Pashkam & Xu, 2017). Moreover, variance-related patterns generalized across orientation, shape and animacy and were detected in visual as well as frontoparietal cortex, indicating that the effects were not confined to a generic attentional or target-monitoring signal. However, attention can modulate not only overall response magnitude but also the fidelity, stability and discriminability of multivoxel representations (Jehee et al., 2011; Ester et al., 2016). We therefore cannot completely exclude an attentional contribution to variance decoding. Future studies should orthogonally manipulate ensemble variance and task relevance, or use an orthogonal task that equates attentional demands across variance conditions, to dissociate variance-related representations from attentional salience.

A complementary interpretation of the DVP involvement, particularly for variance coding, derives from the established role of the parietal cortex in numerical cognition. The IPS is known to represent numerosity, proportions, and non-symbolic fractions (Piazza et al., 2004, 2007; Jacob & Nieder, 2009; Nieder et al., 2013). Because ensemble variance was defined by the relative proportions of category members within an ensemble, variance representations may partly recruit neural mechanisms involved in the non-symbolic encoding of fractional quantities. From this perspective, the present findings extend previous demonstrations of fraction-like coding in the parietal cortex beyond simple dot arrays, showing that proportional information can also be extracted from higher-level ensemble properties such as orientation, shape, and animacy. Notably, however, ROI analyses also revealed feature-general variance representations within the VVP. This suggests that proportion-like information may be represented across multiple visual pathways, potentially linking quantitative coding with the visual feature representations from which ensemble statistics are computed. More broadly, this interpretation strengthens proposed links between ensemble perception and numerical cognition (Chong & Evans, 2011; Corbett et al., 2023). It also raises the intriguing possibility that summary-statistical representations, which are present in preverbal infants (Zosh, Halberda, & Feigenson, 2011) and non-human species (Imura et al., 2017; Lee et al., 2026), may provide early building blocks for the later development of numerical cognition.

Having considered the broader implications of these findings for the neural architecture and domain-generality of ensemble perception, several additional points merit discussion. First, our results suggest that ensemble representations can arise even when participants are not intentionally oriented to identify the predominant class or the variance of the ensemble, underscoring the automaticity and efficiency of ensemble perception (Parkes et al., 2001; Elosegi et al., 2024, 2025). Consistent with previous work (Tark et al., 2021), ensemble mean information was reliably decoded within the early visual cortex despite being behaviourally irrelevant. Moreover, in line with behavioural evidence (Utochkin & Vostrikov, 2019), we observed that some ROIs could represent different ensemble statistics in parallel. An important question for future work is to understand the representational mechanisms that allow multiple ensemble statistics to be encoded simultaneously within the same regions.

Second, the overlap between clusters identified by ensemble-mean regression (Supplementary figure 3) and predominant-class decoding (Figure 3) supports the use of predominant-class judgments as a proxy for average feature representations. This is consistent with our previous behavioural findings, in which participants appeared to discriminate the predominant ensemble class by relying on the average lifelikeness of individual items (Elosegi et al., 2024, 2025). The present neural results therefore provide converging evidence that predominant-class decoding captures ensemble mean information, rather than merely categorical or decision-related signals.

Third, the present findings have implications for computational models of ensemble perception. Population-coding models of ensemble perception and visual crowding have successfully accounted for averaging performance for relatively simple visual features, including orientation, size, colour, and motion direction (Freeman & Simoncelli, 2011; Utochkin et al., 2024). However, such models typically lack the representational depth needed to explain ensemble coding for higher-level features such as shape and animacy (Whitney & Yamanashi Leib, 2018). Our results suggest that future models will need to combine population-level summary mechanisms with hierarchical visual representations capable of encoding increasingly abstract object properties.

In conclusion, these findings suggest that ensemble perception is not supported by a single, domain-general summary mechanism. Instead, the visual system appears to compute different ensemble statistics (i.e., mean and variance) through partially distinct but interacting neural pathways. Ensemble mean representations preserve the structure of the visual hierarchy, remaining closely tied to the features from which they are extracted, whereas ensemble variance is represented more abstractly and generalizes across visual domains. This distinction suggests that feature-specific and feature-invariant ensemble codes coexist within the visual cortex, allowing the visual system to retain information about both the content and structure of complex visual scenes. More broadly, our findings indicate that summary-statistical representations are a fundamental component of visual processing, rather than an auxiliary strategy recruited only when individual-object representations exceed working-memory capacity..

## Acknowledgement

PE was supported by a Basque Government PREDOC grant. DS received support from the Basque Government (BERC 2022–2025), the Spanish Ministry of Economy and Competitiveness (Severo Ochoa Programme, CEX2020-001010-S), and project PID2023-149267NB-I00. YX was supported by the US National Institutes of Health grant R01EY030854. Funders had no role in the study design, data collection, analysis, decision to publish, or manuscript preparation.

## Author contributions

PE, DS, and YX conceptualized and designed the study. PE collected the data and conducted the data analysis in collaboration with ST & NM. PE drafted the manuscript, and DS and YX contributed to writing and critical revisions. All authors reviewed and approved the final version of the manuscript.

