## Supplementary Material for "Neural Representations of Ensemble Mean and Variance Across Visual Features"

### 6. Supplementary Materials

#### 6.1. Validation of CNN-based lifelikeness calculation

We analyzed data from two independent behavioral experiments involving a total of 60 human participants who viewed 12-item ensembles composed of grayscale object images from the Konkle Lab animacy stimulus set (120 images, identical to those used in the current study). Ensembles were arranged in circular formations around fixation and varied in their animacy ratios. In Experiment 1, ratios included 7:5, 8:4, and 9:3 (as well as their symmetric counterparts: 5:7, 4:8, and 3:9), while Experiment 2 included ratios of 6:6, 7:5, 8:4, 9:3, and 10:2 (and their symmetric pairs). On each trial, participants performed a 2-alternative-forced-choice task in which they discriminated the predominant class of the ensemble (animate vs. inanimate), with no time constraints.

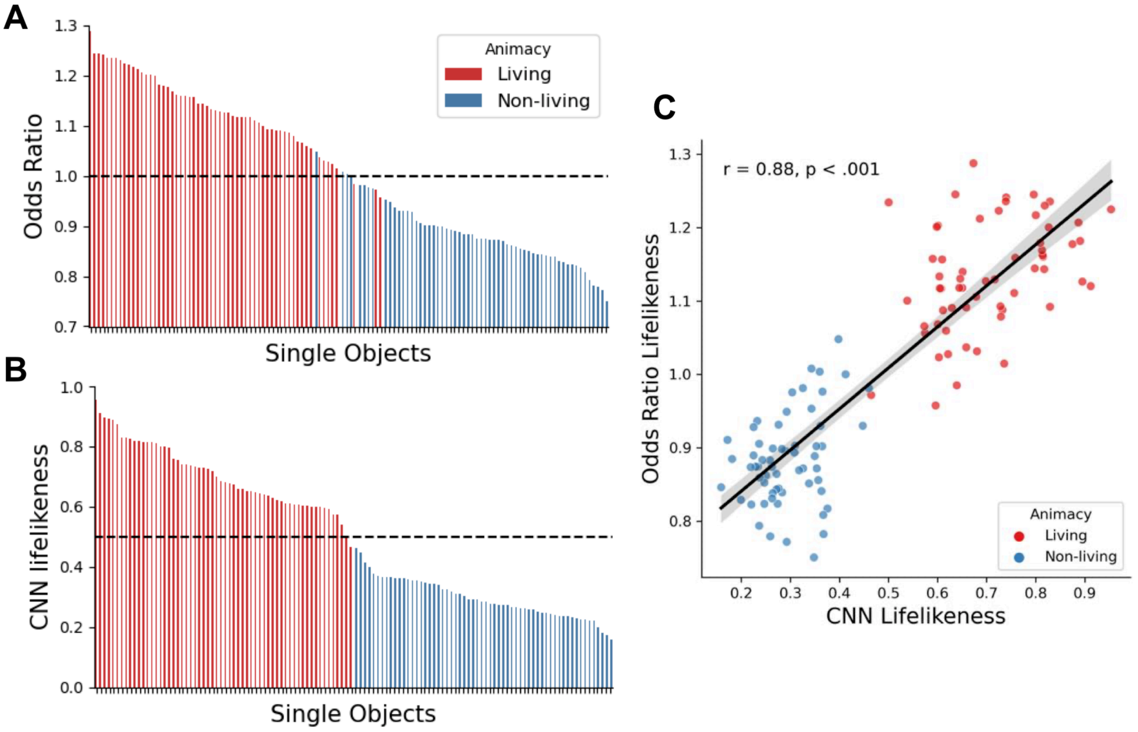


***Supplementary Figure 1 | Correspondence between human- and CNN-derived lifelikeness estimates for single objects.*** **(A)** Odds ratios from a logistic regression predicting human ensemble responses based on the presence of each individual object. Values greater than 1 indicate that the presence of the object increases the likelihood of an ensemble being judged as predominantly animate. Bars are colored by the object's animacy category. **(B)** CNN-derived lifelikeness estimates for the same 120 objects, plotted in descending order and colored by animacy category. **(C)** Correlation between human-based odds ratio lifelikeness scores and CNN-derived lifelikeness scores across the 120 objects. A strong positive correlation was observed ($r = 0.88$, $p < .001$), suggesting that CNNs capture core aspects of human lifelikeness representations at the single-object level.

Experiment 1 manipulated ensemble presentation time and eccentricity on a trial-by-trial basis, resulting in average values of 217 ms and 4.6° visual angle, respectively. In Experiment 2, both parameters were fixed at 150 ms and 4.6°. Each participant completed 1,000 trials, yielding 60,000 trials in total.

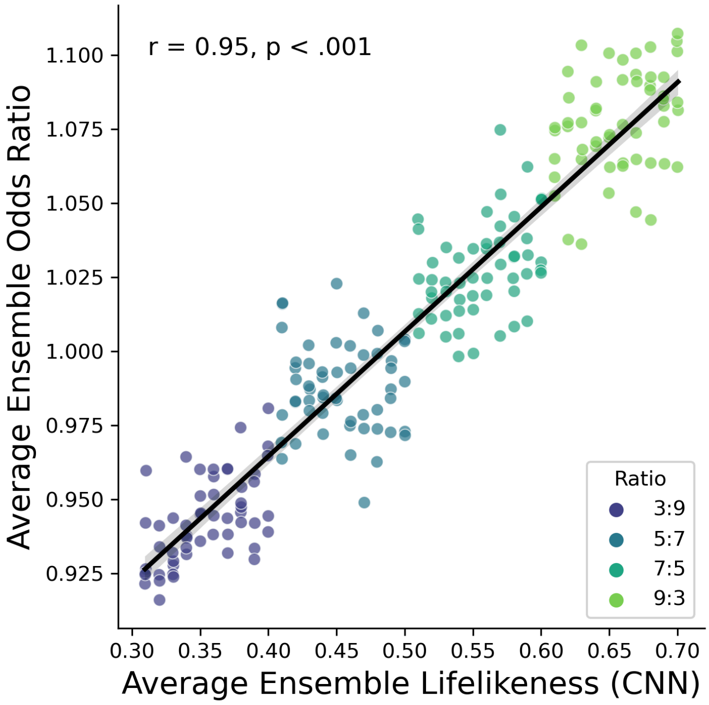


***Supplementary Figure 2 | Correlation between human- and CNN-derived lifelikeness estimates for object ensembles.*** Each point represents one of the 216 ensembles presented in the fMRI study. The X-axis depicts the average CNN-derived lifelikeness scores across the 12 objects in each ensemble. The Y-axis represents the average human-derived odds ratios for the same ensembles, reflecting the likelihood that an ensemble was judged as predominantly animate. Points are color-coded by the ensemble's animacy ratio. A strong positive correlation was observed between human and CNN ensemble scores ($r = 0.95$, $p < .001$), indicating high correspondence between computational and behavioral measures of ensemble lifelikeness.

To assess how the presence of individual objects influenced responses, we constructed a binary matrix with 60,000 rows (trials) and 120 columns (stimuli). Each row indicated whether a specific object was present (1) or absent (0) in a given trial. Since each ensemble contained 12 items selected at random, each object appeared in roughly 9-10% of trials. We trained a logistic regression classifier to predict participants' responses (1 = predominantly animate; 0 = predominantly inanimate) based on this binary input. Using a leave-one-participant-out cross-validation approach, classifiers trained on N–1 participants and tested on the held-out individual achieved a mean ROC-AUC of 0.66 ($p < .001$, non-parametric bootstrap t-test), indicating, as expected, that ensemble composition significantly predicted ensemble judgments.

To quantify the influence of individual items, in each cross-validation fold we extracted odds ratios from the trained classifiers as a measure of feature importance, and then we averaged the resulting odds ratios across folds (see Supplementary Figure 1A). In this context, an odds ratio greater than 1 indicates that the presence of a particular object increases the likelihood of the ensemble being judged as predominantly animate—independently of the other items—due to the random sampling of stimuli. We interpret that if an object increases the probability of perceiving an ensemble as predominantly animate it has to be in virtue of its underlying *lifelikeness* representation. Therefore, we employed the odd ratios as a measure of human lifelikeness values (see Supplementary Figure 1A) to validate our computationally derived lifelikeness estimates (see Supplementary Figure 1B).

An initial visual inspection suggested that human- and CNN-derived lifelikeness scores followed a similar distribution, which was confirmed by a statistically significant Pearson correlation of 0.88 ($p < .001$; see Supplementary Figure 1C). We then computed ensemble-level lifelikeness scores by averaging human-derived odds ratios across the 12 objects shown in each ensemble and correlated these with the corresponding CNN-derived ensemble scores. Notably, the correlation between the two measures increased to 0.95 ($p < .001$; see Supplementary Figure 2). This increase is likely due to the cancellation of random noise across multiple object samples—a well-established phenomenon in psychometrics known as “increased reliability via aggregation” (Lord & Novick, 1968; Rushton, Brainerd, Pressley, 1983), a principled which has been proposed to increase the signal to noise ratio of ensemble representations (Ariely, 2001; Alvarez, 2011). These results suggest that CNN-derived lifelikeness scores provide a robust approximation of human ensemble lifelikeness representations—comparable in reliability to human lifelikeness ratings for single objects, which are commonly used in prior studies (e.g., Leib et al., 2016).

#### 6.2. Whole-brain Searchlight predominant class regression results

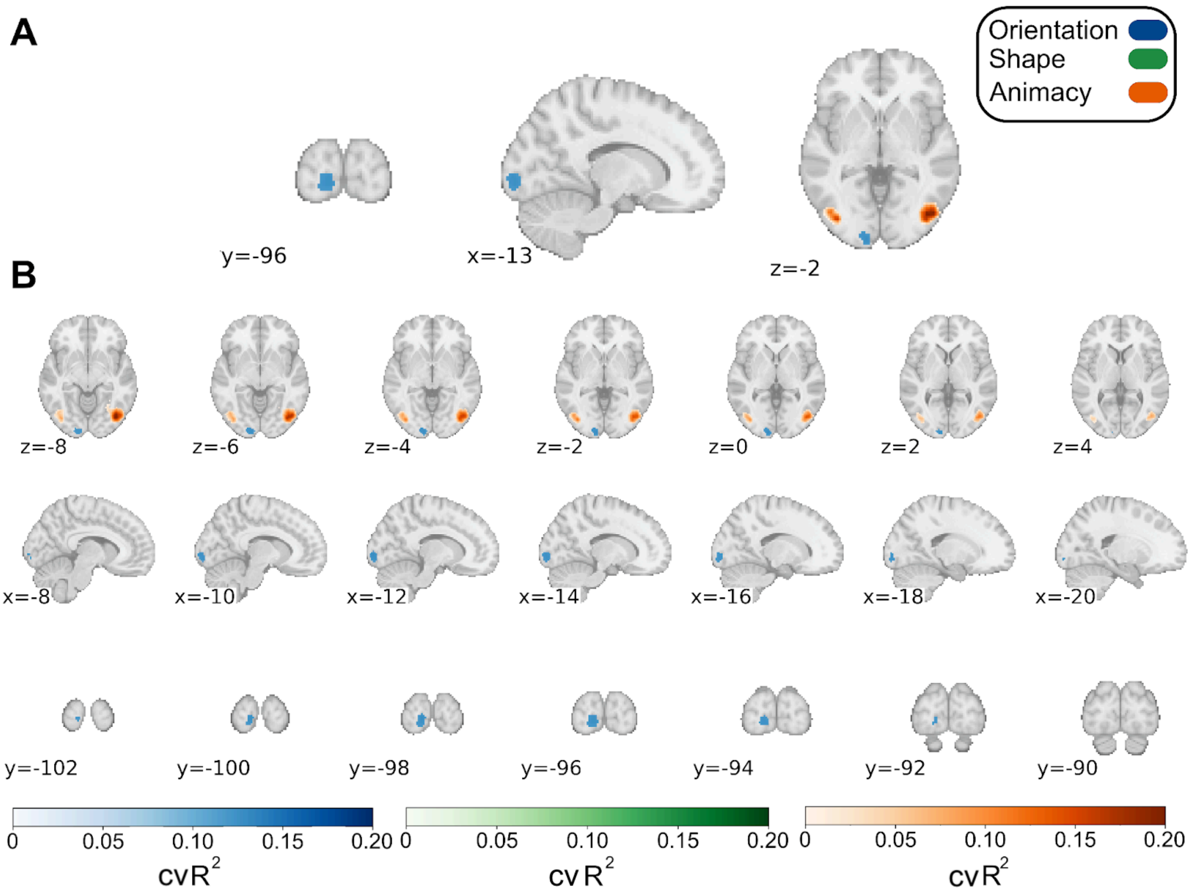


***Supplementary Figure 3 | Whole-Brain Searchlight Regression of Ensemble Mean Across Visual Features.*** The heatmap indicates clusters where cross-validated $R^2$ scores were statistically significantly greater than chance (i.e., $cvR^2$ = 0), determined using threshold-free cluster enhancement (*p* < .05) for each stimulus condition independently. **A)** highlights clusters with the highest $cvR^2$ values across axial, sagittal, and coronal planes. **B)** provides detailed views of cluster locations across the brain, with rows ordered by axial, sagittal, and coronal planes, respectively. All figures are presented in the neurological view (i.e., the left side of the brain appears on the left side of the image).

#### 6.3. Statistical analysis of ROI-based MVPA results

***Supplementary Table 1***
**Group-level paired-samples t-test results for predominant class decoding**

| Decoding | Stim | ROI | $X_{ROC\text{-}AUC}$ | $X_{ROC\text{-}AUC\text{-}Chance}$ | $Perm_{within}$ | t(23) | $p_{Bonf}$ | d | $BF_{10}$ |
|---|---|---|---|---|---|---|---|---|---|
| Within | Ani | DMask | 0.607 | 0.495 | 17/24 | 8.61 | .001 | 1.76 | > 1000 |
| Within | Ani | Fusi | 0.661 | 0.513 | 21/24 | 8.18 | .001 | 1.67 | > 1000 |
| Within | Ani | IPG | 0.606 | 0.502 | 17/24 | 5.84 | <.001 | 1.19 | > 1000 |
| Within | Ani | ITG | 0.59 | 0.495 | 16/24 | 5.63 | <.001 | 1.15 | > 1000 |
| Within | Ani | LatOcc | 0.667 | 0.498 | 21/24 | 10.34 | <.001 | 2.11 | > 1000 |
| Within | Ani | Ling | 0.59 | 0.505 | 16/24 | 5.81 | <.001 | 1.19 | > 1000 |
| Within | Ani | PeriC | 0.512 | 0.496 | 2/24 | 1.28 | 1.000 | 0.26 | 0.78 |
| Within | Ani | SPG | 0.581 | 0.5 | 16/24 | 6.47 | <.001 | 1.32 | > 1000 |
| Within | Ani | VMask | 0.683 | 0.518 | 20/24 | 7.46 | <.001 | 1.52 | > 1000 |
| Within | Ori | DMask | 0.517 | 0.493 | 5/24 | 1.64 | 1.000 | 0.34 | 1.30 |
| Within | Ori | Fusi | 0.527 | 0.494 | 8/24 | 1.87 | 1.000 | 0.38 | 1.83 |
| Within | Ori | IPG | 0.503 | 0.505 | 7/24 | -0.12 | 1.000 | -0.02 | 0.20 |
| Within | Ori | ITG | 0.499 | 0.486 | 2/24 | 1.06 | 1.000 | 0.22 | 0.60 |
| Within | Ori | LatOcc | 0.625 | 0.517 | 18/24 | 5.59 | <.001 | 1.14 | > 1000 |
| Within | Ori | Ling | 0.569 | 0.499 | 15/24 | 5.52 | <.001 | 1.13 | > 1000 |
| Within | Ori | PeriC | 0.576 | 0.505 | 15/24 | 4.35 | .006 | 0.89 | 249.79 |
| Within | Ori | SPG | 0.528 | 0.496 | 8/24 | 1.71 | 1.000 | 0.35 | 1.44 |
| Within | Ori | VMask | 0.609 | 0.516 | 16/24 | 6.07 | <.001 | 1.24 | > 1000 |
| Within | Shp | DMask | 0.58 | 0.495 | 11/24 | 4.37 | .006 | 0.89 | 265.61 |
| Within | Shp | Fusi | 0.557 | 0.496 | 12/24 | 5.03 | .001 | 1.03 | > 1000 |
| Within | Shp | IPG | 0.568 | 0.499 | 8/24 | 4.06 | .013 | 0.83 | 132.07 |
| Within | Shp | ITG | 0.538 | 0.502 | 7/24 | 2.08 | 1.000 | 0.42 | 2.57 |
| Within | Shp | LatOcc | 0.66 | 0.511 | 20/24 | 9.47 | <.001 | 1.93 | > 1000 |

| | | | | | | | | | |
|---|---|---|---|---|---|---|---|---|---|
| Within | Shp | Ling | 0.581 | 0.496 | 15/24 | 5.82 | <.001 | 1.19 | > 1000 |
| Within | Shp | PeriC | 0.587 | 0.51 | 14/24 | 3.86 | .021 | 0.79 | 86.50 |
| Within | Shp | SPG | 0.583 | 0.488 | 12/24 | 6.14 | <.001 | 1.25 | > 1000 |
| Within | Shp | VMask | 0.645 | 0.495 | 20/24 | 9.86 | <.001 | 2.01 | > 1000 |

*Notes and Abbreviations.* $X_{ROC\text{–}AUC}$ = mean decoding performance; $X_{ROC\text{–}AUC\text{–}Chance}$ = Mean of simulated chance-level performance based on permutations; $Perm_{within}$ = fraction of participants with significantly better than chance decoding according to within-subjects permutation test; *d* = Cohen's d effect size; $BF_{10}$ = Bayes Factor as evidence in favor of the alternative hypothesis. Within-decoding (*Within*); animacy (*Anm*); orientation (*Ori*); shape (*Shp*). ROIs included pericalcarine cortex (*PeriC*), lingual gyrus (*Ling*), lateral occipital cortex (*LatOcc*), fusiform gyrus (*Fusi*), inferior temporal gyrus (*ITG*), inferior parietal gyrus (*IPG*), and superior parietal gyrus (*SPG*). The ventral mask (*VMask*) comprised PeriC, Ling, LatOcc, Fusi, and ITG. The dorsal mask (*DMask*) comprised IPG and SPG.

***Supplementary Table 2***

**Group-level paired-samples t-tests and within-subject permutation test results for ensemble variance decoding and cross-decoding**

| Decoding | Stim | ROI | $X_{ROC\text{-}AUC}$ | $X_{ROC\text{-}AUC\text{-}Chance}$ | $Perm_{within}$ | t(23) | $p_{Bonf.}$ | d | $BF_{10}$ |
|---|---|---|---|---|---|---|---|---|---|
| Cross | Ani | DMask | 0.575 | 0.512 | 13/24 | 6.2 | <.001 | 1.27 | > 1000 |
| Cross | Ani | Fusi | 0.539 | 0.51 | 5/24 | 2.88 | .230 | 0.59 | 11.02 |
| Cross | Ani | IPG | 0.573 | 0.51 | 12/24 | 5.77 | <.001 | 1.18 | > 1000 |
| Cross | Ani | ITG | 0.542 | 0.494 | 7/24 | 4.02 | .014 | 0.82 | 121.98 |
| Cross | Ani | LatOcc | 0.531 | 0.497 | 4/24 | 2.86 | .238 | 0.58 | 10.71 |
| Cross | Ani | Ling | 0.531 | 0.485 | 5/24 | 3.87 | .021 | 0.79 | 87.85 |
| Cross | Ani | PeriC | 0.525 | 0.497 | 2/24 | 2.92 | .209 | 0.6 | 11.93 |
| Cross | Ani | SPG | 0.566 | 0.49 | 10/24 | 4.28 | .008 | 0.87 | 213.50 |
| Cross | Ani | VMask | 0.56 | 0.509 | 9/24 | 5.34 | <.001 | 1.09 | > 1000 |
| Cross | Ori | DMask | 0.623 | 0.501 | 16/24 | 7.59 | <.001 | 1.55 | > 1000 |
| Cross | Ori | Fusi | 0.557 | 0.505 | 7/24 | 2.57 | .459 | 0.53 | 6.16 |
| Cross | Ori | IPG | 0.624 | 0.492 | 16/24 | 9.76 | <.001 | 1.99 | > 1000 |
| Cross | Ori | ITG | 0.555 | 0.502 | 5/24 | 2.88 | .231 | 0.59 | 10.98 |
| Cross | Ori | LatOcc | 0.583 | 0.494 | 13/24 | 6.02 | <.001 | 1.23 | > 1000 |
| Cross | Ori | Ling | 0.556 | 0.518 | 8/24 | 2.21 | 1.000 | 0.45 | 3.20 |
| Cross | Ori | PeriC | 0.538 | 0.494 | 6/24 | 3.47 | .056 | 0.71 | 36.92 |

| | | | | | | | | | |
|---|---|---|---|---|---|---|---|---|---|
| Cross | Ori | SPG | 0.599 | 0.491 | 13/24 | 6.73 | <.001 | 1.37 | > 1000 |
| Cross | Ori | VMask | 0.604 | 0.488 | 14/24 | 6.55 | <.001 | 1.34 | > 1000 |
| Cross | Shp | DMask | 0.611 | 0.487 | 18/24 | 8.16 | <.001 | 1.67 | > 1000 |
| Cross | Shp | Fusi | 0.564 | 0.499 | 9/24 | 5.09 | .001 | 1.04 | > 1000 |
| Cross | Shp | IPG | 0.601 | 0.503 | 14/24 | 6.66 | <.001 | 1.36 | > 1000 |
| Cross | Shp | ITG | 0.562 | 0.502 | 9/24 | 4.0 | .015 | 0.82 | 116.23 |
| Cross | Shp | LatOcc | 0.576 | 0.489 | 11/24 | 7.87 | <.001 | 1.61 | > 1000 |
| Cross | Shp | Ling | 0.549 | 0.5 | 7/24 | 3.85 | .022 | 0.79 | 84.34 |
| Cross | Shp | PeriC | 0.51 | 0.513 | 0/24 | -0.35 | 1.000 | -0.07 | 0.17 |
| Cross | Shp | SPG | 0.598 | 0.507 | 13/24 | 7.58 | <.001 | 1.55 | > 1000 |
| Cross | Shp | VMask | 0.589 | 0.488 | 13/24 | 6.21 | <.001 | 1.27 | > 1000 |
| Within | Ani | DMask | 0.558 | 0.497 | 12/24 | 4.4 | .006 | 0.9 | 278.97 |
| Within | Ani | Fusi | 0.533 | 0.482 | 6/24 | 3.74 | .029 | 0.76 | 66.66 |
| Within | Ani | IPG | 0.553 | 0.492 | 11/24 | 5.08 | .001 | 1.04 | > 1000 |
| Within | Ani | ITG | 0.549 | 0.491 | 6/24 | 3.87 | .021 | 0.79 | 88.36 |
| Within | Ani | LatOcc | 0.532 | 0.509 | 6/24 | 1.76 | 1.000 | 0.36 | 1.53 |
| Within | Ani | Ling | 0.532 | 0.488 | 7/24 | 3.62 | .039 | 0.74 | 50.58 |
| Within | Ani | PeriC | 0.516 | 0.497 | 2/24 | 1.48 | 1.000 | 0.3 | 1.02 |
| Within | Ani | SPG | 0.565 | 0.499 | 10/24 | 5.04 | .001 | 1.03 | > 1000 |
| Within | Ani | VMask | 0.551 | 0.508 | 9/24 | 3.5 | .052 | 0.72 | 39.75 |
| Within | Ori | DMask | 0.69 | 0.513 | 23/24 | 9.55 | <.001 | 1.95 | > 1000 |
| Within | Ori | Fusi | 0.617 | 0.486 | 17/24 | 5.83 | <.001 | 1.19 | > 1000 |
| Within | Ori | IPG | 0.67 | 0.514 | 23/24 | 7.05 | <.001 | 1.44 | > 1000 |
| Within | Ori | ITG | 0.626 | 0.485 | 17/24 | 8.07 | <.001 | 1.65 | > 1000 |
| Within | Ori | LatOcc | 0.641 | 0.516 | 19/24 | 6.02 | <.001 | 1.23 | > 1000 |
| Within | Ori | Ling | 0.614 | 0.505 | 18/24 | 5.48 | <.001 | 1.12 | > 1000 |
| Within | Ori | PeriC | 0.577 | 0.491 | 13/24 | 5.02 | .001 | 1.03 | > 1000 |
| Within | Ori | SPG | 0.681 | 0.503 | 24/24 | 10.51 | <.001 | 2.14 | > 1000 |

| | | | | | | | | | |
|---|---|---|---|---|---|---|---|---|---|
| Within | Ori | VMask | 0.665 | 0.491 | 22/24 | 7.8 | <.001 | 1.59 | > 1000 |
| Within | Shp | DMask | 0.622 | 0.507 | 18/24 | 7.45 | <.001 | 1.52 | > 1000 |
| Within | Shp | Fusi | 0.592 | 0.487 | 15/24 | 8.39 | <.001 | 1.71 | > 1000 |
| Within | Shp | IPG | 0.616 | 0.507 | 19/24 | 6.7 | <.001 | 1.37 | > 1000 |
| Within | Shp | ITG | 0.593 | 0.493 | 15/24 | 6.32 | <.001 | 1.29 | > 1000 |
| Within | Shp | LatOcc | 0.6 | 0.5 | 17/24 | 5.48 | <.001 | 1.12 | > 1000 |
| Within | Shp | Ling | 0.552 | 0.497 | 11/24 | 4.01 | .015 | 0.82 | 118.37 |
| Within | Shp | PeriC | 0.537 | 0.506 | 10/24 | 2.41 | .651 | 0.49 | 4.60 |
| Within | Shp | SPG | 0.614 | 0.5 | 17/24 | 6.75 | .001 | 1.38 | > 1000 |
| Within | Shp | VMask | 0.615 | 0.497 | 18/24 | 7.37 | <.001 | 1.5 | > 1000 |

*Notes and Abbreviations.* $X_{ROC-AUC}$ = mean decoding performance; $X_{ROC-AUC-Chance}$ = Mean of simulated chance-level performance based on permutations; $Perm_{within}$ = fraction of participants with significantly better than chance decoding according to within-subjects permutation test; *d* = Cohen's d effect size; $BF_{10}$ = Bayes Factor as evidence in favor of the alternative hypothesis. Within-decoding (*Within*); Cross-decoding (*Cross*); animacy (*Anm*); orientation (*Ori*); shape (*Shp*). ROIs included pericalcarine cortex (*PeriC*), lingual gyrus (*Ling*), lateral occipital cortex (*LatOcc*), fusiform gyrus (*Fusi*), inferior temporal gyrus (*ITG*), inferior parietal gyrus (*IPG*), and superior parietal gyrus (*SPG*). The ventral mask (*VMask*) comprised PeriC, Ling, LatOcc, Fusi, and ITG. The dorsal mask (*DMask*) comprised IPG and SPG.
